# Species boundaries structure competitive interactions among honeybee gut bacteria

**DOI:** 10.64898/2026.07.28.741209

**Authors:** Silvia Brochet, Germán Bonilla-Rosso, Florent Mazel, Philipp Engel

## Abstract

Bacterial species often harbor extensive strain-level diversity, raising the question of whether strains, rather than species, are the relevant ecological units. We quantified pairwise interactions among 12 strains from four prevalent *Lactobacillus* species of the honeybee gut microbiota using gnotobiotic bees. Microbiota-depleted bees were colonized with individual strains and all 66 pairwise combinations, with strain abundances measured by colony counts and strain-resolved amplicon sequencing. Negative interactions predominated, with most significant interactions involving mutual inhibition. Within-species interactions were stronger and more asymmetric than between-species interactions, producing less even community compositions. Across repeated cycles of colonization in microbiota-depleted bees, three of four within-species pairs lost one strain, whereas all four between-species pairs persisted. These results indicate that closely related strains experience greater niche overlap and stronger competition than strains from different species, supporting bacterial species as ecologically differentiated units while highlighting the importance of strain-specific traits in shaping community assembly and stability.

## Introduction

Similar bacteria often thrive under comparable environmental conditions because they share physiological traits and have similar energy and nutrient requirements [1–3]. As a result, closely related species, and even strains of the same species, frequently co-occur in the same environment [4–10].

According to the bacterial species concept proposed by Cohan (2002)[11], bacterial diversification (that is, the buildup of new bacterial lineages) is driven by adaptation to distinct ecological niches, with species representing ecologically cohesive units. By contrast, strains within a species are generally assumed to occupy largely overlapping niches [12], implying stronger ecological similarity. Under this framework, competition is expected to be stronger among conspecifics (strains of the same species) than among allospecifics (strains of different species) [13, 14]. However, the widespread co-occurrence of both conspecific and allospecific strains in natural communities challenges this expectation [15]. Moreover, extensive intraspecific variation in gene content suggests that conspecific strains could exploit different resources or conditions, occupying distinct functional niches in the same environment [16–22]. Together, these observations suggest that ecological differentiation in bacteria operates not only between species but also within them, with conspecific strains potentially constituting discrete ecological units comparable to species. Under this view, patterns of bacterial coexistence may be governed at the strain level rather than the species level. Indeed, a recent study of pitcher plant microbiomes showed that the dynamics of closely related strains (differing by as little as 100 base pairs) can markedly differ within the same community, indicating that strains of the same species respond differently to ecological conditions [10]. Similarly, macroecological properties of the human gut microbiome, such as stability, appear to be determined at the strain level [23].

These findings raise an important question: Do strains within the same species interact in fundamentally different ways than strains from different species, and can such differences explain observed patterns of coexistence?

While bacterial interactions have been experimentally studied at both the strain and species levels [24–30], systematic comparisons across these levels remain limited. Moreover, such interactions are frequently studied *in vitro*, outside the natural environments in which these microbes occur, limiting the generalizability of the findings.

Honeybees provide a powerful model to address this question. Their gut microbiota is dominated by a small number of bacterial genera, each comprising several closely related species that harbor substantial diversity at the strain level [19, 31–34]. Importantly, strains from most species are readily culturable, and microbiota-depleted bees can be generated and experimentally colonized with defined synthetic communities assembled from these isolates [35, 36].

One of the most prevalent genera in the honey bee gut is *Lactobacillus* (formerly referred to as Firm-5) [31, 33, 37]. Isolates from this group are closely related and typically fall into four to five distinct species [19, 38], which likely diverged from a common ancestor associated with honeybees. Each of these species harbors substantial strain-level diversity. Both species and, to some extent, strains within *Lactobacillus* co-occur in conventional bees [19].

Moreover, previous experiments demonstrated that four strains, each representing a distinct species, can stably coexist in gnotobiotic bees when provided with a pollen and sugar water diet, but not when provided with sugar water alone [25]. These results indicate that niche partitioning of pollen-derived nutrients underlies their coexistence. However, it remains unclear to what extent these interactions are representative of the broader diversity within each species, and whether interactions among strains within species differ fundamentally from those observed between species.

Here, we selected three strains for each of four honeybee-specific *Lactobacillus* species (n=12 strains) and colonized microbiota-depleted bees with pairs of these strains in all possible combinations (n=66 combinations). This allowed us to systematically measure the type and strength of interactions between all possible between-species and within-species interactions among the selected strains. We then selected eight pairs to study their ability to stably co-exist over several generation of bees. Our results show that negative interactions dominate within the bee gut microbiome, and that their strength is greater within than between bacterial species. Therefore, strains belonging to the same species were more likely to be partially displaced by one another. Over multiple bee generations, conspecific strain pairs were less likely to co-exist than pairs from different species, suggesting that stronger negative interactions can destabilize long-term co-existence.

## Results

### Colonization of gnotobiotic bees with all pairwise combinations of 12 strains from four *Lactobacillus* species

To assess differences between species-level and strain-level interactions within the bee gut, we selected three strains from each of the four *Lactobacillus* species mentioned above (in total 12 strains, see Materials and Methods, **Supplementary Table 1**). Pairwise average nucleotide identity (ANI) between strains of the same species varied between 96.03-97.85%, while ANI among strains of different species was <85%, indicating clear differences in divergence between within- and between-species pairs (**Supplementary Figure 1**). We colonized microbiota-depleted (MD) bees with each of the 12 strains separately as well as in all 66 possible pairwise combinations (**Figure 1A, Supplementary Table 2**). Ten days post-colonization, we dissected the distal hindgut (rectum) of 3-5 bees per treatment (in total 332 bees) and quantified the total abundance of each community member by combining CFU plating with amplicon sequencing of a house-keeping gene with strain-level sequence variation (*mutM*). This approach enables discrimination among the selected strains (**Figure 1B**), and was established in our previous work [25]. The inocula used for the colonization of MD bees were analyzed using the same procedure as the homogenized bee guts to evaluate the potential impact of inoculum strain ratios on colonization success.

**Figure 1.**
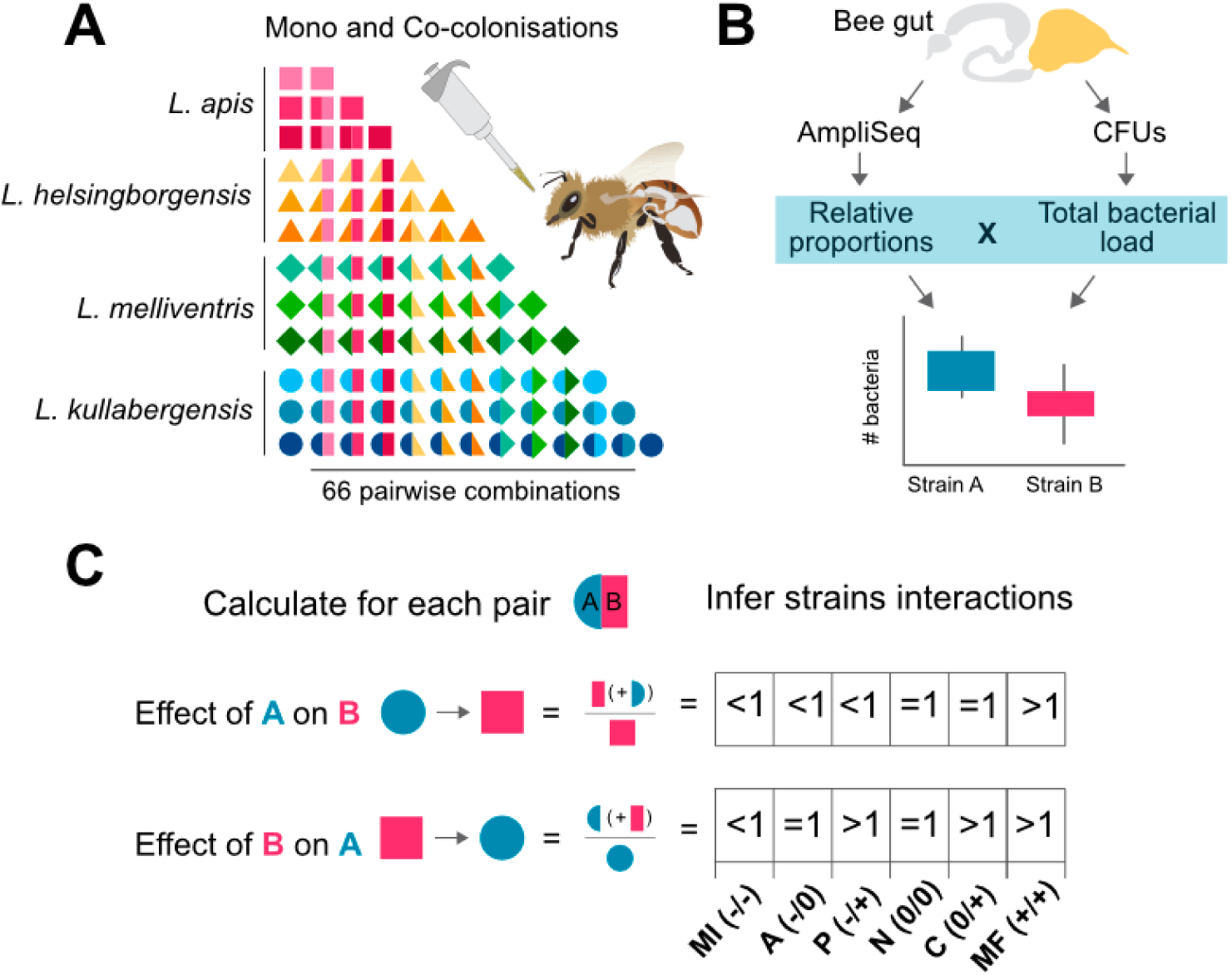
Experimental design and analytical framework for systematically testing pairwise strain interactions in gnotobiotic bees. **(A)** Twelve strains (different colors) from four prevalent *Lactobacillus* species (indicated by different shapes) of the bee gut microbiota were selected to analyze species-vs strain-level interactions in the bee gut: *Lactobacillus apis*, *Lactobacillus helsingborgensis*, *Lactobacillus melliventris*, *Lactobacillus kullabergensis*. Microbiota-depleted (MD) bees were colonized with each of the 12 strains separately as well as in all 66 possible pairwise combinations (n=3-5 per treatment). **(B)** Ten days post-colonization the rectum of the gnotobiotic bees was subjected to CFUs plating and amplicon sequencing to determine the absolute abundance of each of the two community members in each strain pair by multiplying the relative proportion of each strain in the community with the total bacterial load. **(C)** To determine the interactions governing each strain pair, the effect of one strain on the other was calculated by computing the ratio between the bacterial loads in the co-colonization versus the mono-colonization condition for each strain. Depending on the reciprocal effects of the two strains on each other, the type of interaction was classified as either mutual inhibition (MI, -/-), amensalism (A, -/0), parasitism (P, -/+), neutralism (N, 0/0), commensalism (C, 0/+), or mutual facilitation (MF, +/+).

CFU counting and amplicon sequencing were successful for all samples (5.0 × 10o–5.0 × 10o CFUs per gut; 1k–55k reads per sample; Supplementary Figure 2). Based on these two measures we determined the limit of detection (LOD, due to limited sequencing depth) for each strain within a given sample (10^4^-10^6^ copies per bee gut, **Supplementary Figure 2**). While most gut samples were dominated by the two inoculum strains, a small fraction of samples (5.4%) had contamination (>10% reads assigned to strains not included in the inoculum, **Supplementary Figure 3**) and were excluded. Although we aimed to colonize all bees with equal proportions of the two strains (see Methods), one strain was often more abundant than the other in the inoculum but these small initial difference had no effect on strain proportions ten days post inoculation (**Supplementary Figure 4**), indicating that interactions with the host, the diet, or the other community members in the gut determined community composition rather than the initial composition of the inoculum. We found that co-colonisation loads were higher than both monoculture loads in 32% of pairs (21/66), a pattern analogous to overyielding in agricultural science and community ecology, while in 30% of pairs (20/66) co-colonisation exceeded only one monoculture load, and in 38% of pairs (25/66) both monoculture loads were higher than the co-colonisation load (**Supplementary Figure 5**).

### Negative interactions dominate among the 12 *Lactobacillus* strains

Both strains of the inoculum were detected in all gut samples after 10 days of colonization suggesting that none of the strains completely outcompeted the other one. To analyze the type of interaction governing each strain pair we used a similar approach as Kehe et al. [39]. In short, we quantified each one-way interaction within a given sample, that is, the effect of strain A on strain B (and vice versa). To do so, we calculated the ratio of the yields of strain B in co-colonization and mono-colonization (see Methods, **Formula 1**). A ratio <1 was classified as a negative (-) effect, a ratio >1 was classified as a positive (+) effect, and a ratio of 1 was classified as no effect (0) of strain A on strain B. The two one-way interactions for each strain pair were then combined, yielding six possible bidirectional interaction types (-/-, -/0, +/-, 0/0, +/0, or +/+; **Figure 1C**). To capture the quantitative aspects of these interactions, we represented them as vectors in polar coordinates, defined by an angle (θ) and a radius (r) (see Materials and Methods, Formula 2; **Figure 2A**), as measures of the asymmetry and the cumulative effect size of the interaction, respectively. In cases of mutual inhibition (−/−), where one strain exerts a substantially stronger inhibitory effect on the other strain than it receives in return, the angle θ will be less negative than when both strains inhibit each other to a similar extent. For example, Ө = −90° indicates mutual inhibition of equal strength, while Ө = −45° indicates amensalism, where one strain has a negative effect on the other while experiencing no effect in return (extreme asymmetry). Similarly, if the combined mutual inhibition is relatively weak (i.e. not much change in total abundance compared to the mono-colonizations), the radius r will be smaller than in a case where the two strains in sum inhibit each other more strongly.

**Figure 2.**
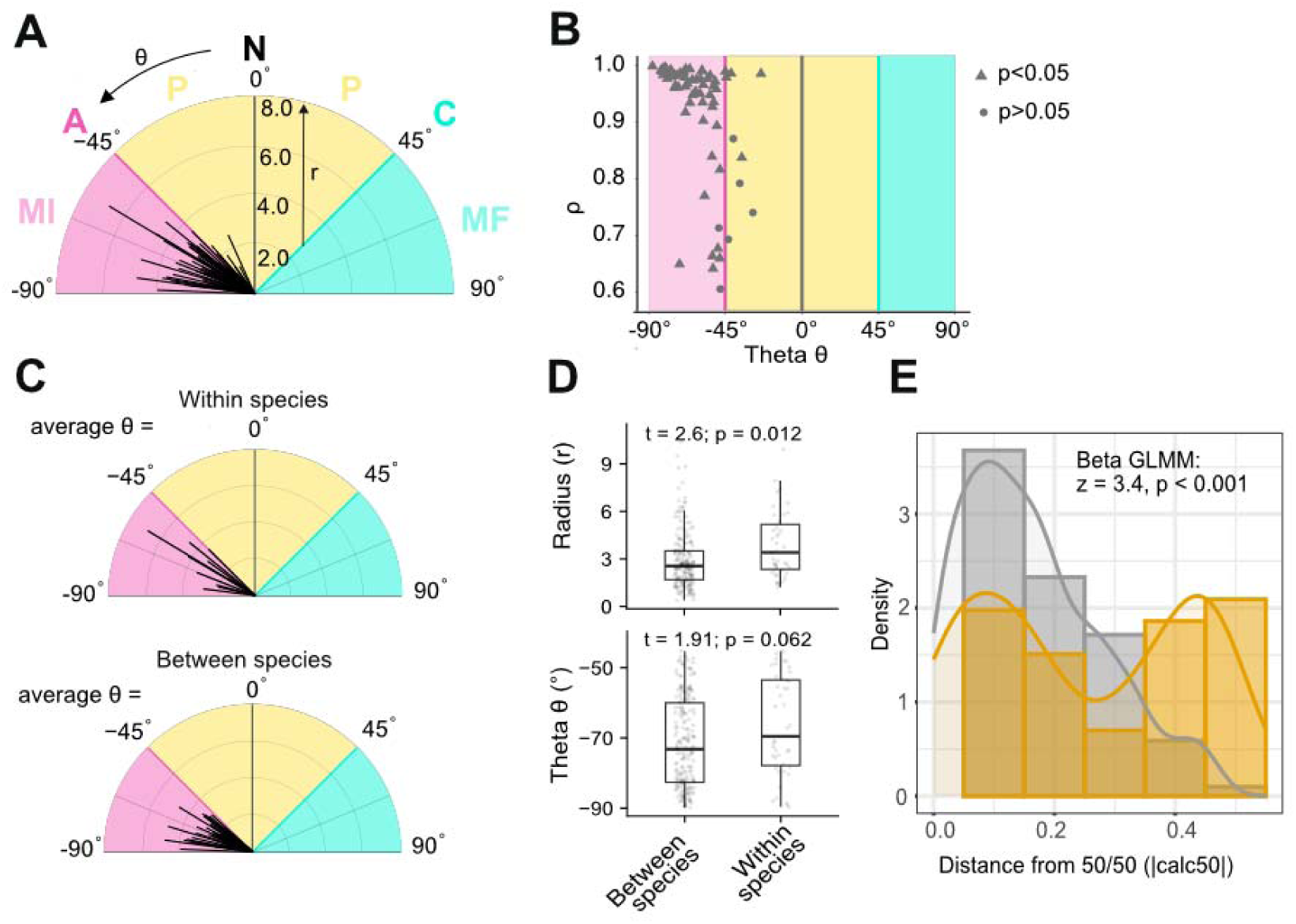
Polar coordinates describing pairwise strain interaction across the 12 *Lactobacillus* strains. (A) Polar plot displaying means of replicates for each strain pair (n=3-5) as vectors of angle θ (type of interaction) and length r (strength of interaction). The different colors represent the different interaction types (mutual inhibition, MI; amensalism, A; parasitism, P; neutralism, N; commensalism, C; mutual facilitation, MF). Note that A, N, and C only exist if the angle θ is exactly, -45°, 0°, and 45°, respectively. (B) Statistical analysis of vectors dispersion between the replicates of each strain pair. Each point represents one pair of strains. r is a measure of the vector’s dispersion between the different replicates of a strain pair. It can range from 0 (high dispersion) to 1 (low dispersion). The significance of r was tested for each vector for each replicate using circular statistics (see Materials and methods). (C) Polar plots displaying means of vector replicates for each strain pair divided into pairs of strains from the same species and different species. (D) Boxplots of competitive interaction strength (radius, top) and directionality (theta, bottom) for within-species and between-species pairs within the competitive quadrant (theta < −45°). Dots show individual replicate values. Statistics from linear mixed models with Pair, Strain A, and Strain B identity as random effects. (E) Distribution of competitive dominance (absolute distance from 50/50 relative abundance) across competitive pairs (theta < −45°), colored by pair type (yellow: within-species; grey: between-species). Histograms and density curves are shown at the replicate level. Statistical comparison by beta mixed model with group-specific dispersion.

To assess whether a given pair exhibits a robust type of interaction, i.e. consistent results across bees colonized with the same pair, we calculated the dispersion of the vectors across these samples (r, 0 = high dispersion, 1 = low dispersion). We then applied circular statistics (see Methods) to identify vectors with significantly low dispersion, indicating consistent and well-explained interactions. For 60 out of 66 pairs, the two strains had a statistically significant interaction type (Pycke’s test, p < 0.05), i.e. their corresponding vectors were concentrated around one preferred direction and were not dispersed around the circle (**Figure 2B**) [40]. Of these significant interactions, 56 were categorized as mutual inhibition and 4 as parasitism (**Figure 2B**). Notably, although most interactions were categorized as mutual inhibition, their θ values varied substantially across pairs, ranging from -24.2° to -87.8°, indicating that interactions were more balanced in some pairs than in others. Likewise, the r values of these interaction vectors also varied substantially across pairs ranging from 1.16 to 7.71 indicating that some pairs had stronger effects on each other than others.

Taken together, we conclude that most of the interactions between the tested *Lactobacillus* strains are negative, but that there is variation in the type and strength of reciprocal inhibitory effect of the two strains on each other.

### Negative interactions within species are more biased that those between species

While negative interactions predominated among most of the tested *Lactobacilli*, their strength and type varied, potentially reflecting underlying phylogenetic or functional divergence between strains within each pair. Focusing on strain pairs showing mutual inhibition (see Methods), we found that pairs from different species (allospecific pairs) exhibited lower *r* values (t=2,6, p<.05, mixed linear models) and a tendency toward lower θ values compared to pairs from the same species (**Figure 2C and 2D**). This effect was also visible (but not statistically significant) when we correlated both the r (interaction strength) and the θ (interaction type) values with both the genetic distance and the differences in the accessory gene content between the strains in each pair (**Supplementary Figure 6**). This pattern indicates stronger and less balanced reciprocal effects in within-species relative to between-species interactions. Consistent with this, we observed differences in the relative abundance of the two community members across strain pairs. Between-species pairs tended to be more centered around the 50/50 distribution while within-species pairs were further apart, with greater variation in strain proportions (mixed generalized model, z=2.7, p<.001, **Figure 2E**). Altogether, these results show subtle but statistically significant differences in how strains within and between species interact with consequences on community composition.

### Strains of different species stably coexist, while strains of the same species tend to exclude each other across multiple bee generations

Although none of the tested strains was able to outcompete any of the other strains in the co-colonization experiment, the disproportional abundances of the two strains in within-species in respect to between-species pairs suggest that strains of the same species may not be able to coexist over longer time across bee generations. To explore this, we serially passaged eight randomly selected strain pairs (four between-species and four within-species pairs) through MD bees for a total of three passages (P0-P2, **Supplementary Figure 7**, **Figure 3A**). After each passage (i.e. after 7 days of colonization), we used the same method as above, i.e. amplicon sequencing in combination with CFUs counting, to determine the absolute abundance of each strain in each pair (**Figure 3B and 3C**).

**Figure 3.**
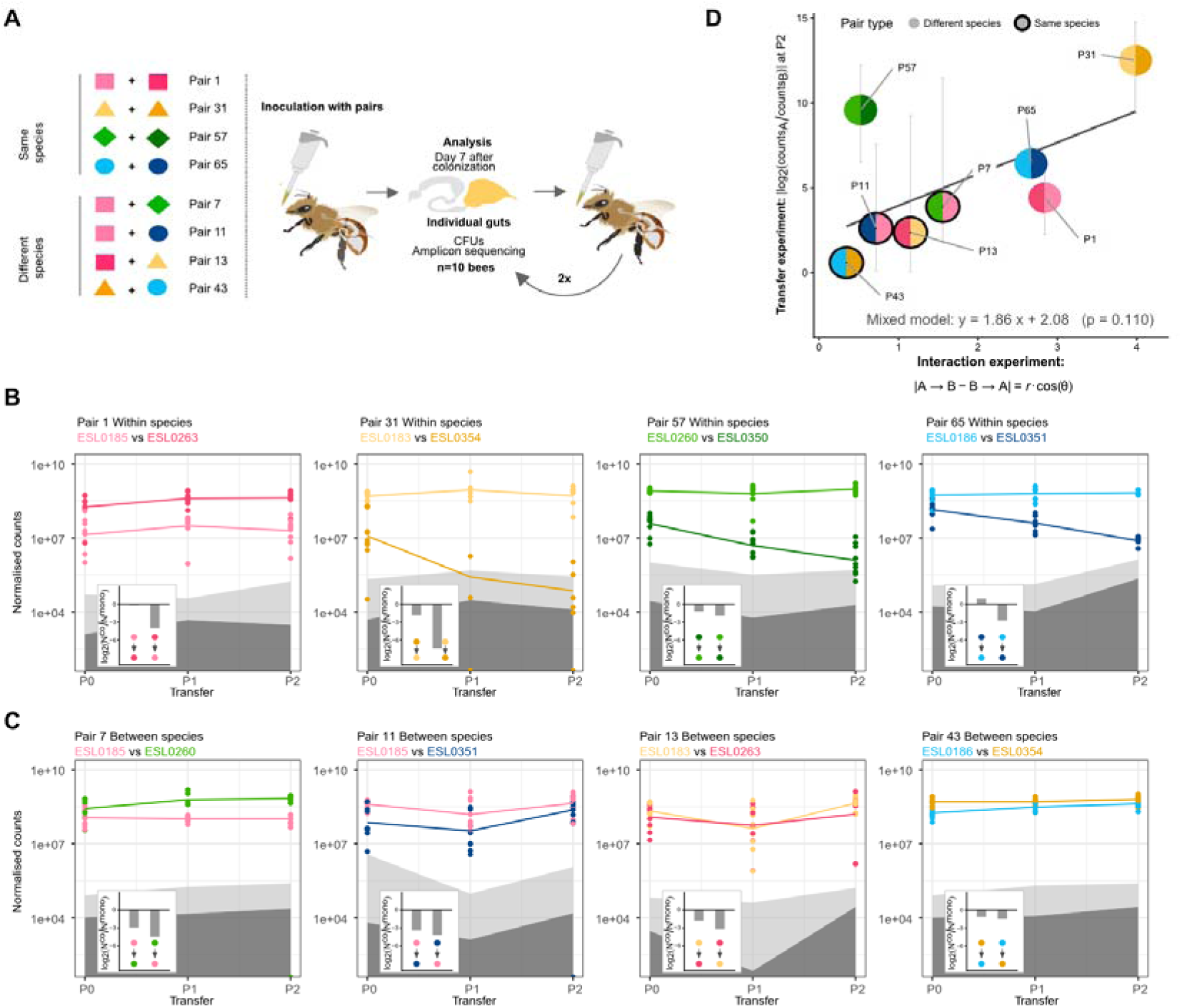
Differential coexistence patterns in conspecific and allospecific strain pairs in gnotobiotic Bee. **(A)** Experimental setup of the in vivo passaging experiment, comprising four pairs of strains from the same species (pairs 1, 31, 57, 65) and four pairs from different species (pairs 7, 11, 13, 43). Microbiota-depleted bees were colonized with the selected strain pairs and seven days post-colonization the rectum was dissected. An aliquot was used for CFUs plating and amplicon sequencing. The remaining gut content of each replicate was pooled to colonize a new batch of microbiota-depleted bees. In total three passages through gnotobiotic bees (P0-P2) were performed. **(B and C)** Bacterial counts per rectum (n=10) for the two strains in each of the eight tested strain pairs across the three passages (P0-P2). Counts were determined by multiplying the total number of CFUs with the relative abundance of each strain in the gut as determined by amplicon sequencing. Grey areas represent the limit of detection: the 95% confidence intervals of the limit of detection are shown in light grey. Insets show the pairwise interaction strengths measured in the independent co-culture experiment, expressed as log□(NC^co / NC^mono), where NC is the normalized count (CFU × relative abundance, Figure 2). Each bar represents one strain’s NC fold-change in co-culture relative to monoculture; colored dots indicate the acting strain (top) and the affected strain (bottom), with the arrow pointing from acting to affected. (**D)** Relationship between interaction asymmetry measured in the pairwise co-culture experiment and strain dominance at the end of the transfer experiment. The x-axis shows the interaction asymmetry metric r·cos(θ), derived from the co-culture interaction space: larger values indicate stronger asymmetry, with one strain more suppressed than the other. The y-axis shows the absolute log□ ratio of normalized counts of Strain A to Strain B at the final transfer (P2), averaged across replicates (error bars: min–max range). The grey line is the fitted relationship from a linear mixed model with a random intercept per pair. The equation and p-value refer to the r·cos(θ) coefficient.

In three out of the four within-species pairs (pairs #31, #57 and #65, see **Supplementary Table 2**), one of the two strains steadily decreased in abundance (**Supplementary Table 3**, negative slope with p <0.05), with two of them reaching the limit of detection after one and two passages, respectively (**Figure 3B**). In contrast, in the fourth pair (pair 1), while the two strains exhibited clear differences in the bacterial loads in the gut, none of them decreased in abundance over the three passages suggesting stable coexistence (**Figure 3B, Supplementary Table 3**, p >0.05). A similar pattern was observed for all four between-species pairs (**Supplementary Table 3**, p >0.05). Both strains were consistently detected across all passages in all four pairs, with neither approaching the limit of detection nor exhibiting a decline in total abundance over successive passages (**Figure 3C**). However, two pairs (pairs #11 and #13, see **Supplementary Table 2 and 4**) exhibited slightly more variation in bacterial abundance over the three passages (especially at P1) than the other two pairs (pair 7 and 43, see **Supplementary Table 2 and 4**) (**Figure 3C**). In summary, these results indicate that only one out of four within-species pairs, but all four between-species pairs can stably coexist over three passages through MD bees (Fisher’s exact test, one-sided, p = 0.071). The co-existence dynamics across the three transfer passages broadly agreed with the results of the interaction experiment (compare main and inset panels of **Figure 3B, 3C,** and **Supplementary Figure 7**). In general, the strain least affected by its partner in the interaction experiment maintained higher and more stable counts across passages. To test this more formally, we combined the two polar-coordinate parameters from the interaction experiment into a single metric, r·cos(θ). The radial distance r captures overall interaction strength, while the angle θ encodes interaction symmetry: at θ = −90°, both strains are equally suppressed and cos(θ) = 0; as θ approaches −45°, the boundary of the competitive quadrant, one strain becomes increasingly unaffected while the other remains suppressed, and cos(θ) increases accordingly. The product r·cos(θ) is therefore large only when competition is simultaneously strong and asymmetric and is proportional to the absolute difference between the two interaction effects (|A→B − B→A|). Pairs with higher r·cos(θ) — predominantly within-species pairs — showed more extreme count ratios at passage P2, consistent with near-exclusion of one strain but this trend was not statistically significant (**Figure 3D**; slope = 1,86, p =0,1). The clearest example is pair #31: ESL0354, heavily suppressed by ESL0183 in the interaction experiment, rapidly fell below the detection limit during the transfer experiment. A notable exception was pair #57, where ESL0350 nearly fell below the detection limit at P2 despite both strains showing similar inhibitory effects on each other in the interaction experiment.

## Discussion

A longstanding question in microbiology is whether species constitute ecologically distinct units [41–43]. If so, pairs of strains within the same species should exhibit greater niche overlap, and consequently stronger negative interactions, than pairs of strains from closely related but distinct species.

Here, we examined this question by measuring *in vivo* pairwise interactions and coexistence among strains of four closely related *Lactobacillus* species of the honey bee gut microbiota. These species belong to a bee-specific clade of *Lactobacillus* and are largely restricted to the gut of honeybees, suggesting diversification into multiple strains and species within this environment [19, 31, 38, 44]. This system provides a valuable model to test whether interactions within and between closely related species differ fundamentally in their native context.

Our results show that negative interactions dominate between the tested strains regardless of whether they are from the same or different species. However, these interactions were, on average, stronger and less symmetric between strains of the same species than between strains of different species, and resulted in less even community compositions (i.e. one strain tend to dominate the other more often when both strains belonged to the same bacterial species). Consistent with this pattern, within-species pairs were more likely than between-species pairs to lose one of the two strains over longer timescales spanning multiple bee generations (transfer experiment). Together, these findings support the idea that strains of the same species interact differently than strains of different species and hence differ in their niche overlap and ecology. Our results help explain diversity patterns observed in the gut of conventional bees where species appear to co-occur within, while strains of the same species tend to segregate across individual hosts [19]. Similar patterns have been observed for the human microbiota [8, 45, 46].

Yet, the differences between interaction metrics of between-species and within-species pairs were modest, and we observed substantial variation in the strength and type of interactions among both types of strain pairs. For example, the four pairs exhibiting parasitic (−/+) interactions included both between-species and within-species pairs. Moreover, one of the four within-species pairs tested in the passaging experiment showed evidence of stable coexistence, whereas in the other three cases, one of the two strains excluded the other. These findings underscore the importance of strain-level identity and sub-species variation in shaping bacterial interactions and their consequences for community assembly [12, 24, 47] For example, strain-resolved surveys of the gut microbiota in conventional honey bees have shown that strain composition deviates significantly from random expectations [6, 19, 48], suggesting that strains might indeed differ in their ecological niches. Moreover, although strain diversity is typically higher between individuals than within a single bee in natural populations, multiple strains have nonetheless been detected within individual hosts [19, 31, 48, 49]. Similar patterns have been reported in other microbial communities, both in host-associated and free-living environments suggesting that community dynamics, and consequently community functions, are at least to some extent driven by strain-level variation [8, 10, 23, 24, 47, 50–52].

A somewhat unexpected finding of our study was that most interactions were mutually negative (mutual inhibition), even among strains from different species. These negative interactions could result from multiple non-mutually processes such as direct antagonism (e.g. via bacterial toxins), competition for shared nutrients or space. We suggest that competition for shared nutrients might at least contribute to some of these negative interactions for two reasons. First, previous work reported strong priority effects among bee gut bacteria: firstcomer strains prevented latercomer strains to establish, even when belonging to different *Lactobacillus* species [53, 54], suggesting that species of the same genus have large niche overlap and small fitness differences. Second, empirical work manipulating host diet shows that the presence of pollen in the bee diet has been shown to increase the total number of bacterial cells a gut can sustain, specifically for the genus *Lactobacillus* [25, 56]. This suggests that the carrying capacity in the bee gut is limited either by the availability of pollen-derived compounds (such as co-factors, nitrogen and carbon sources) in the rectum or by niche space, which may expand in the presence of pollen (for example through increased surface area available for colonization) [57]. Theory predicts that if competition for nutrient (partial niche overlap) is the only process driving negative interactions and fitness differences are small, we should observe overyielding (i.e., total co-colonization loads exceeding both mono-colonization loads). However, we only observed overyielding in about 1/3 of the pairs, suggesting that direct antagonism (e.g. via bacteriocin), large fitness differences among strains or niche modification (e.g. via pH changes) also underly these negative interactions [55].

A previous study based on pairwise interaction tests between 67 bacterial species reported that both metabolic similarity and evolutionary relatedness can predict the presence of negative interactions [58]. Although we examined a much narrower phylogenetic range of bacteria, our results are broadly consistent with this finding: more closely related taxa (here, strains within the same *Lactobacillus* species) exhibited, on average, stronger negative interactions than more distantly related taxa (here, strains belonging to different *Lactobacillus* species). However, we did not detect a statistically significant relationship between phylogenetic distance or accessory gene content similarity and either the strength or type of interaction. This likely reflects limitations of our experimental design, as the strain pairs we tested were either very closely related or very distantly related, leaving intermediate distances underrepresented.

Nevertheless, prior comparative genomic analyses have revealed differences in metabolic potential among strains of the four species included in our study [38]. In particular, variation was observed in the number and types of genes involved in carbohydrate metabolism (e.g., COG category ‘G’, PTS transporters, and glycoside hydrolases) between species [59]. Consistent with these findings, GC-MS-based metabolomic analyses of culture supernatants of strains of the four *Lactobacillus* species examined here showed differences in the utilization of sugars and sugar acids [25]. Together, these results indicate substantial metabolic diversity among strains and suggest that strains from different bacterial species have less metabolic niche overlap than strains within the same species.

A key question is whether species co-existence, community assembly and stability can be predicted from these pairwise interaction measurements [60, 61]. We use the term coexistence here with an important caveat. Stable coexistence is formally defined by mutual invasibility (Chesson 2000; Grainger et al. 2019): each species must exhibit a positive per-capita growth rate when rare while the competitor is at equilibrium density. Our experiments do not constitute a formal test of this criterion because we initiated all competitions at approximately equal proportions (∼50:50) and tracked outcomes over three passages (∼0–2 generations). We therefore measure short-term competitive persistence, the absence of rapid extinction, rather than stable coexistence in the strict theoretical sense. Our results suggest that identifying negative interactions alone is insufficient to predict competitive exclusion at the timescale we measured it: five of the eight tested pairs coexisted stably over multiple passages, despite competing for the same carrying capacity in the bee gut. Moreover, although strain pairs exhibiting stronger negative interactions on each other were also more likely to exclude each other in the passaging experiment (mostly within-species pairs), this trend was not significant. This may reflect limited statistical power, as we had relatively few replicates per strain pair, and substantial variability in interaction vectors among individual bees within the same pair. It is also known that other stabilizing mechanism can mediate co-existence in the presence of competition, such as environmental fluctuations [61–63].

Interestingly, one of the strains (ESL0263) in the only within-species pair which showed evidence of coexistence, was also the only strain in our previous work that did not experience priority effects of other strains [53]. This strain was able to establish in the bee gut even when introduced three days after pre-colonization with a 12-member community composed of *Bifidobacterium*, *Lactobacillus*, and *Bombilactobacillus*. Future studies will be needed to characterize the functional traits that confer this strain’s apparent competitive advantage.

Environmental context can strongly influence the outcome of strain interactions and community assembly [64]. A key strength of our study design is that interactions were tested in the native environment of these bacteria (i.e. in the honey bee gut) in the presence of the natural host diet (pollen). However, the gnotobiotic bees lacked any gut microbiota members other than the two *Lactobacillus* strains tested. The presence of the native bee gut community could modulate niche overlap and, consequently, interaction outcomes between focal strains. For example, other community members may open new metabolic niches due to cross-feeding interactions or further restrict them due to nutrient niche partitioning and competition for space, changing strain interaction and coexistence patterns. Therefore, to what extent strain interactions measured in simple settings can be extrapolated to complex communities remains to be tested in the bee study system. Several studies have documented higher-order interactions determining community composition and dynamics across systems and environments [65, 66], while others suggest that negative interaction networks may still be reasonably inferred from pairwise measurements [67]. The bee gut model provides an excellent system to test how different community backgrounds shape interaction strength between conspecific and allospecific strain pairs in future studies, with implications for our fundamental understanding of microbial ecology and the functional relevance of strain-level diversity in this ecologically important pollinator host.

## Materials and Methods

### Culturing of bacterial strains

We used 12 *Lactobacillus* strains, three of each of the following four species: *Lactobacillus apis*, *Lactobacillus helsingborgensis*, *Lactobacillus melliventris* and *Lactobacillus kullabergensis*, see **Supplementary Table 1**). All strains were inoculated precultured from glycerol stocks stored at −80°C onto solid De Man – Rogosa – Sharpe agar (MRSA), supplemented with 2% w/v fructose and 0.2% w/v L-cysteine-HCl. Plates were incubated for three days in anaerobic conditions at 34°C. Then, single colonies were picked and inoculated into liquid carbohydrate-free MRS medium (cfMRS [68]) supplemented with 4% glucose (w/v), 4% fructose (w/v), and 1% L-cysteine-HCl (w/v) and incubated at 34°C in anaerobic conditions without shaking.

### Colonization of microbiota-depleted bees

Microbiota-depleted bees were obtained from colonies of Apis mellifera carnica located at the University of Lausanne following the procedure described in [36]. Bacterial colonization stocks were prepared from overnight cultures by washing the bacteria in 1xPBS, diluting them to an OD_600_ = 1, mixing them in all pairwise combinations (see **Supplementary Table 2**), and storing them in 25% glycerol at −80°C until further use. Colonization stocks were diluted ten times in a 1:1 mixture of 1xPBS and sugar water (50% sucrose solution, w/v) and 5 μL were fed to each microbiota-depleted bee 1 days after eclosion using a pipette. Bees were kept on a sugar water + pollen diet and food was provided ad libitum. Ten days post-colonization, five rectums were dissected and homogenized in 1xPBS. An aliquot of each homogenized rectum was used for CFU plating to enumerate the total bacterial load and for amplicon sequencing to obtain the relative abundance of each community member.

For the eight strain pairs that were passaged through microbiota-depleted bees, a total of ten rectums were dissected seven days post-colonization and homogenized in 1xPBS. An aliquot of each homogenized rectum was used to plate CFUs and to perform amplicon sequencing. The remaining gut homogenates were pooled together and stored at −80°C until further use. At the day of colonization, a frozen aliquot of the pooled gut homogenate was thawed, diluted ten times in a 1:1 mixture of 1xPBS and sugar water (50% sucrose solution, w/v), and fed to newly emerged microbiota-depleted bee as described above. This was repeated for a total of three serial passages.

### Assessing colonization levels using amplicon sequencing

The relative abundance of each strain across all experiments was obtained using a previously established amplicon sequencing approach based on a 199-bp long fragment of the gene *mutT,* which allowed us to discriminate the 12 strains from each other [59]. In brief, a crude cell lysate was prepared by mixing 5 μL of gut homogenate with 50 μL of lysis solution, containing 45 μL of lysis buffer (10 mM Tris-HCl, 1 mM EDTA, 0.1% Triton, pH 8), 2.5 μL of lysozyme (20 mg/ml, Fluka), and 2.5 μL of Proteinase K solution (10 mg/ml, Roth). The samples were incubated for 10 min at 37°C, for 20 min at 55 °C, and for 10 min at 95 °C, followed by a short spin before preparing the PCR (1 min, 1500 rpm). The gene fragment was then amplified from the crude cell lysate following a two-step PCR protocol. To prepare the first PCR mix, 5 μL of cell lysate were mixed with 12.5 μL of GoTaq Colorless Master Mix (Promega), 1 μL of forward and reverse primer (5 μM) and 5.5 μL of Nuclease-free Water (Promega). The PCR was performed as follows: initial denaturation (95°C – 3 min), 30 times denaturation-annealing-extension (95°C – 30 s, 64°C – 30 s, 72°C – 30 s), final extension (72 °C – 5 min). To purify the amplicons, 15 μL of PCR product were mixed with 5 μL of a 5X Exo-SAP solution (15% Shrimp Alkaline Phosphatase – 1000 U/ ml – NEB, 10% Exonuclease I – 20,000 U/ ml – NEB, 45% glycerol 80% and 30% dH2O), and incubated for 30 min at 37°C and for 15 min at 80°C. To prepare the second PCR mix, 5 μL of purified PCR products were mixed with the same reagents as before. The PCR program was the same as above with the exception that the annealing temperature was set to 60°C and the denaturation-annealing-extension steps were repeated for only eight times.

To prepare the sequencing of the amplicons, DNA concentrations were measured using Quant-iT PicoGreen for dsDNA (Invitrogen). Each sample was adjusted to a DNA concentration of 0.5 ng/μL and 5 μL of each sample were pooled together. The pooled sample was loaded on a 0.9% agarose gel and gel-purified using the QIAquick Gel Extraction Kit (Qiagen) following the manufacturer’s instructions. Then the purified DNA was prepared for sequencing following the Illumina Miniseq guidelines and loaded on a Illumina MiniSeq Mid Output Reagent Cartridge using the correspondent MiniSeq flow cell. Illumina reads were demultiplexed based on the sample-specific barcodes and quality-filtered using Trimmomatic (LEADING:28 TRAILING: 29 SLIDING WINDOW:4:15 MINLEN:90) [69]. Each forward and reverse read pair was assembled using PEAR (-m 290 n 284 j 4 -q 26 v 10 -b 33) [70]. The resulting sequences were then assigned to each strain using a custom-made script and a closed reference database containing the gene variants of minT found in the genomes of the 12 Lactobacillus strains. The proportion of each strain in a sample was calculated as the number of reads assigned to that strain divided by the total number of assigned reads. The absolute abundance of each strain was then determined by multiplying the total CFUs measured for the sample by this proportion.

### Quantification of pairwise strain interactions

To quantify bidirectional interactions between the two strains of each pair we followed the approach established by [39] and outlined in **Supplementary Figure 8**. In short, we calculated the log□ ratio of the abundance of strain A during co-colonization with strain B and its abundance during mono-colonization, and vice versa (**Formula 1**).

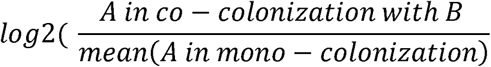

**Formula 1:** Quantification of the effect of strain B on strain A based on differences between mono- and co-colonization levels.

A positive log□ ratio indicates that strain B has a positive effect on strain A (+), whereas a negative log□ ratio indicates a negative effect (-). A log□ ratio of 0 indicates that strain B has no effect on strain A (0). By combining each strain’s effect on the other, we determined the bidirectional interaction type: mutual inhibition (MI, -/-), amensalism (A, -/0), parasitism (P, -/+), neutralism (N, 0/0), commensalism (C, 0/+), or mutual facilitation (MF, +/+). The bidirectional log2 ratios of each replicate for all strain pair were plotted as cartesian coordinates to visualize the reciprocal effect of the two strains on each other (**Supplementary Figure 8**). Following Kehe et al., coordinates were flipped where necessary so that Strain A always denotes the more inhibited partner (A_on_B ≤ B_on_A), ensuring a consistent directional convention across all pairs. These coordinates were then converted to polar coordinates to obtain a quantitative measure of the type (angle θ) and the strength of each reciprocal interaction (radius r) using the following (**Formula 2**).

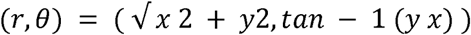

**Formula 2:** Conversion of cartesian coordinates into polar coordinates.

More specifically, every coordinate is converted from a point of coordinates x and y into a vector whose length or radius (r) describes the distance of the vector from the center and whose angle (θ) describes the position of the vector on the plane (**Supplementary Figure 8**). To align the angle θ with the interaction-type convention described above, θ was rotated by +45° relative to the standard polar angle such that perfectly symmetric mutual inhibition (A_on_B = B_on_A) corresponds to θ = −90°, and the amensalism boundary (A_on_B = 0) corresponds to θ = −45°. The radius r is a measure of the cumulative strength of the bidirectional interaction. The angle θ indicates not only the type of the interaction (mutual inhibition or -/-: -45° − -90°; ammensalism or -/0: -45°; parasitism or -/+: 45° − -45°; commensalism or 0/1: 45°; mutual facilitation or +/+: 45° − 90°) but also how symmetric this interactions are between the two strains (**Supplementary Figure 5**). To determine within which pairs strains were interacting significantly, a ρ measure of the dispersion of the replicate vectors for each pair was calculated. Pycke’s test was used to determine which pairs replicate vectors were concordant and thus indicating that the measured interaction was significant [40]. For pairs in figure 3, we characterised the strength and asymmetry of each competitive interaction using the metric r·cos(θ), derived from the polar-coordinate representation of the pair-averaged interaction effects. The radial distance r captures overall interaction strength. The angle θ encodes symmetry: at θ = −90°, both strains are equally suppressed (cos(θ) = 0); as θ approaches −45° — the boundary of the competitive quadrant — the interaction becomes increasingly one-sided, and cos(θ) increases toward its maximum. The metric r·cos(θ) is therefore proportional to the absolute difference in the two interaction effects, |A→B − B→A|, and is large only when competition is both strong and asymmetric. This metric was used as the predictor in Fig. 3C, where it is plotted against the absolute log□ count ratio between the two strains at passage P2, tested using a linear mixed model with pair identity as a random effect.

### Interaction experiment analysis

We focused on strain showing mutual inhibition (“competitive”). To do so, first, we identified competitive pairs at the pair level. A pair was classified as competitive if (i) its mean interaction angle (θ) across replicates fell below −45°, placing it in the competitive quadrant of the interaction space (i.e., both strains were on average suppressed relative to monoculture), and (ii) its replicates showed significant directional clustering around that mean, as assessed by the Pycke test of uniformity on a circle (p < 0.05). This yielded 56 competitive pairs (10 within-species, 46 between-species). Second, within these 56 selected pairs, individual replicates whose interaction angle exceeded −45° (i.e., strayed outside the competitive quadrant) were excluded from the replicate-level analyses.

To test whether competitive interaction strength (radius) and directionality (theta) differed between same-SDP and different-SDP pairs, we fitted linear mixed models (R package lmerTest) on replicate-level data restricted to the competitive quadrant (theta < −45°), with pair type as fixed effect and pair, strain A, and strain B identities as random effects to account for non-independence among pairs sharing a strain. To focus analyses of interaction asymmetry and dominance on pairs that showed genuine competitive interactions, we applied a two-step selection procedure to the 66 co-culture pairs.

Competitive dominance was quantified as the absolute distance from 50/50 relative abundance (|0.5 − relative abundance|) per replicate. To compare same-SDP and different-SDP pairs, we fitted a beta-distributed generalized linear mixed model (R package glmmTMB; logit link) with pair type as fixed effect, pair, strain A, and strain B identities as random effect, and a group-specific dispersion submodel (dispformula = ∼Status) to account for heterogeneous variance between groups. Values were rescaled from (0, 0.5) to (0, 1).

### Transfer experiment analysis

To assess whether strain abundances changed directionally over the course of the pairwise transfer experiment, we fitted a linear model to the log□□-transformed normalised counts (amplicon-based relative abundance scaled by CFU/mL) of each focal strain as a function of transfer number (P0 = 0, P1 = 1, P2 = 2): log□□(normalised counts) ∼ transfer. Observations with zero normalised counts were excluded. The slope estimate and its associated p-value were extracted from each model (Supplementary Table 3). A strain was classified as showing a significant declining trend if the slope was negative and p < 0.05 (linear model t-test). A pair was classified as “declining” if at least one of its two focal strains showed a significant negative slope across transfers. The association between pair type (same vs. different SDP) and declining trend (yes/no) was tested using a one-sided Fisher’s exact test (alternative: same-SDP pairs more likely to decline).

### Code and data availability

Raw sequencing data are archived in NCBI SRA under BioProject PRJNA1497736: https://www.ncbi.nlm.nih.gov/bioproject/?term=PRJNA1497736. The. The code and all other data types needed to reproduce the analysis (CFU counts, metadata, reference sequences) can be found on Github: https://github.com/FloMazel/Bee-Gut-Bacteria-Interactions-Brochet_et_al.

## Acknowledgements

We thank the Lausanne Genomics Technology Facility (GTF) team at the University of Lausanne for performing amplicon sequencing for this study. This work was supported by the NCCR Microbiomes, a National Centre of Competence in Research funded by the Swiss National Science Foundation (SNSF, grant number 180575 and 225148, to P.E.), the SNSF Consolidator grant GLOBEE (213860, to P.E.), the ERC Starting grant MicroBeeOme (714804, to P.E.), and the SNSF starting grant 226295 (to F.M).

## Supplementary Tables and Figures

**Supplementary Table 1.** List of bacterial strains used in this study.

|  |  |  |  |
| --- | --- | --- | --- |
| *Lactobacillus apis* | ESL0263 | GCA_003692845.1 | [37] |
|  | ESL0185 | GCA_003150935.1 | [35] |
|  | ESL0353; Hma11 | GCA_000970735.1 | [69] |
| *Lactobacillus helsingborgensis* | ESL0183 | GCA_003173695.1 | [35] |
|  | wkb8 | GCA_000761135.1 | [70] |
|  | ESL0354; Bma5 | GCA_000970855.1 | [69] |
| *Lactobacillus melliventris* | ESL0184 | GCA_003202825.1 | [35] |
|  | ESL0260 | GCA_003692935.1 | [37] |
|  | ESL0350; Hma8 | GCA_000970775.1 | [69] |
| *Lactobacillus kullabergensis* | ESL0186 | GCA_003151025.1 | [35] |
|  | ESL0261 | GCA_003693025.1 | [37] |
|  | ESL0351; Biut2 | GCA_000967195.1 | [69] |

**Supplementary Table 2.** List of all tested combinations between the 12 selected *Lactobacillus* strains.

| Combination ID | Strain 1 | Strain 2 |
| --- | --- | --- |
| 1 | ESL0185 | ESL0263 |
| 2 | ESL0185 | ESL0353 |
| 3 | ESL0185 | ESL0183 |
| 4 | ESL0185 | ESL0354 |
| 5 | ESL0185 | wkb8 |
| 6 | ESL0185 | ESL0184 |
| 7 | ESL0185 | ESL0260 |
| 8 | ESL0185 | ESL0350 |
| 9 | ESL0185 | ESL0186 |
| 10 | ESL0185 | ESL0261 |
| 11 | ESL0185 | ESL0351 |
| 12 | ESL0263 | ESL0353 |
| 13 | ESL0263 | ESL0183 |
| 14 | ESL0263 | ESL0354 |
| 15 | ESL0263 | wkb8 |
| 16 | ESL0263 | ESL0184 |
| 17 | ESL0263 | ESL0260 |
| 18 | ESL0263 | ESL0350 |
| 19 | ESL0263 | ESL0186 |
| 20 | ESL0263 | ESL0261 |
| 21 | ESL0263 | ESL0351 |
| 22 | ESL0353 | ESL0183 |
| 23 | ESL0353 | ESL0354 |
| 24 | ESL0353 | wkb8 |
| 25 | ESL0353 | ESL0184 |
| 26 | ESL0353 | ESL0260 |
| 27 | ESL0353 | ESL0350 |
| 28 | ESL0353 | ESL0186 |
| 29 | ESL0353 | ESL0261 |
| 30 | ESL0353 | ESL0351 |
| 31 | ESL0183 | ESL0354 |
| 32 | ESL0183 | wkb8 |
| 33 | ESL0183 | ESL0184 |
| 34 | ESL0183 | ESL0260 |
| 35 | ESL0183 | ESL0350 |
| 36 | ESL0183 | ESL0186 |
| 37 | ESL0183 | ESL0261 |
| 38 | ESL0183 | ESL0351 |
| 39 | ESL0354 | wkb8 |
| 40 | ESL0354 | ESL0184 |
| 41 | ESL0354 | ESL0260 |
| 42 | ESL0354 | ESL0350 |
| 43 | ESL0354 | ESL0186 |
| 44 | ESL0354 | ESL0261 |
| 45 | ESL0354 | ESL0351 |
| 46 | wkb8 | ESL0184 |
| 47 | wkb8 | ESL0260 |
| 48 | wkb8 | ESL0350 |
| 49 | wkb8 | ESL0186 |
| 50 | wkb8 | ESL0261 |
| 51 | wkb8 | ESL0351 |
| 52 | ESL0184 | ESL0260 |
| 53 | ESL0184 | ESL0350 |
| 54 | ESL0184 | ESL0186 |
| 55 | ESL0184 | ESL0261 |
| 56 | ESL0184 | ESL0351 |
| 57 | ESL0260 | ESL0350 |
| 58 | ESL0260 | ESL0186 |
| 59 | ESL0260 | ESL0261 |
| 60 | ESL0260 | ESL0351 |
| 61 | ESL0350 | ESL0186 |
| 62 | ESL0350 | ESL0261 |
| 63 | ESL0350 | ESL0351 |
| 64 | ESL0186 | ESL0261 |
| 65 | ESL0186 | ESL0351 |
| 66 | ESL0261 | ESL0351 |

**Supplementary Table 3.** Linear model estimates for each focal strain across the pairwise transfer experiment.

| Interaction | Pair | Strain | <sup>1</sup>Slope_est | P_value | Sig_neg_slope |
| --- | --- | --- | --- | --- | --- |
| Between species | 7 | ESL0185 | -0.021 | 0.754 |  |
| Between species | 7 | ESL0260 | 0.206 | 0.014 |  |
| Between species | 11 | ESL0185 | 0.019 | 0.857 |  |
| Between species | 11 | ESL0351 | 0.231 | 0.162 |  |
| Between species | 13 | ESL0183 | 0.177 | 0.283 |  |
| Between species | 13 | ESL0263 | 0.067 | 0.739 |  |
| Between species | 43 | ESL0186 | 0.19 | < 0.001 |  |
| Between species | 43 | ESL0354 | 0.05 | 0.238 |  |
| Within species | 1 | ESL0185 | 0.078 | 0.568 |  |
| Within species | 1 | ESL0263 | 0.182 | 0.063 |  |
| Within species | 31 | ESL0183 | 0.016 | 0.858 |  |
| Within species | 31 | ESL0354 | -1.135 | 0.001 | * |
| Within species | 57 | ESL0260 | 0.033 | 0.602 |  |
| Within species | 57 | ESL0350 | -0.746 | < 0.001 | * |
| Within species | 65 | ESL0186 | 0.047 | 0.448 |  |
| Within species | 65 | ESL0351 | -0.614 | < 0.001 | * |

<sup>1</sup>For each pair and focal strain, a linear model was fitted to log<sub>10</sub>-transformed normalised counts (amplicon-based relative abundance × CFU/mL) as a function of transfer number (P0–P2). Observations with zero normalised counts were excluded. Slope estimate and associated p-value are reported. An asterisk (\*) indicates a significant negative slope (slope < 0, p < 0.05), used to classify pairs as declining for the Fisher's exact test (Fig. 3).

**Supplementary Table 4.**
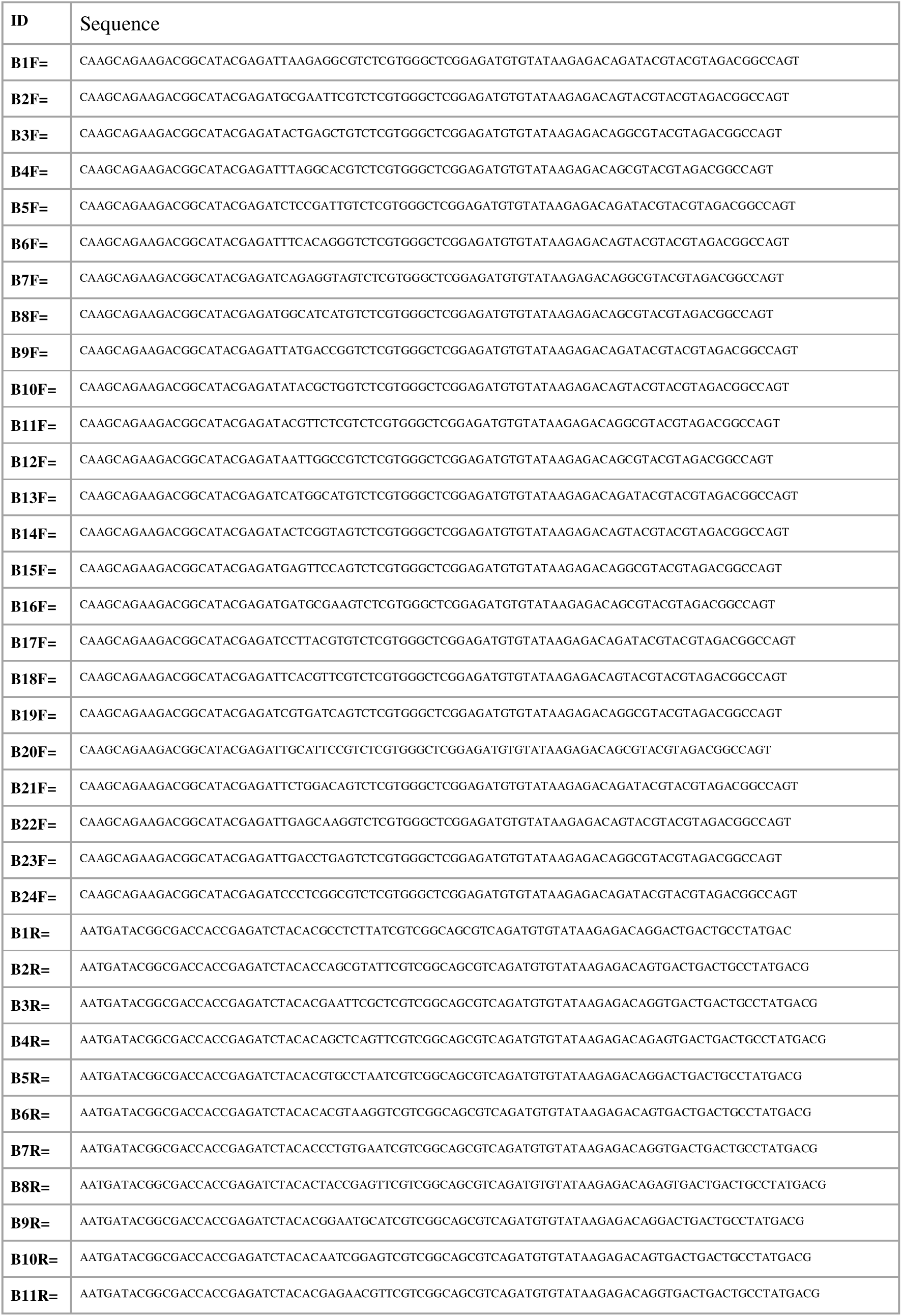
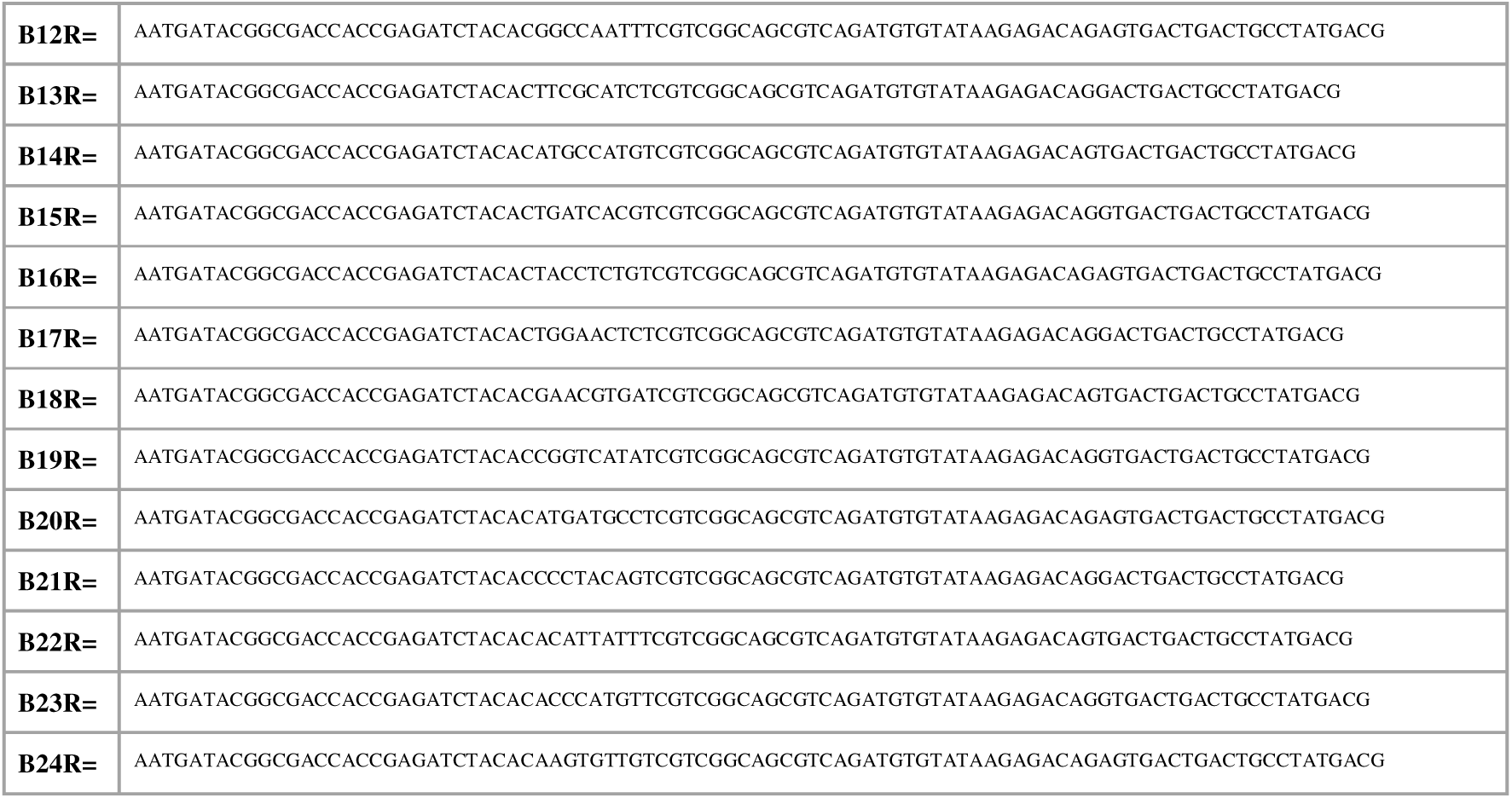
List of barcoded primers used in this study.

## Supplementary Figures

**Supplementary Figure 1.**
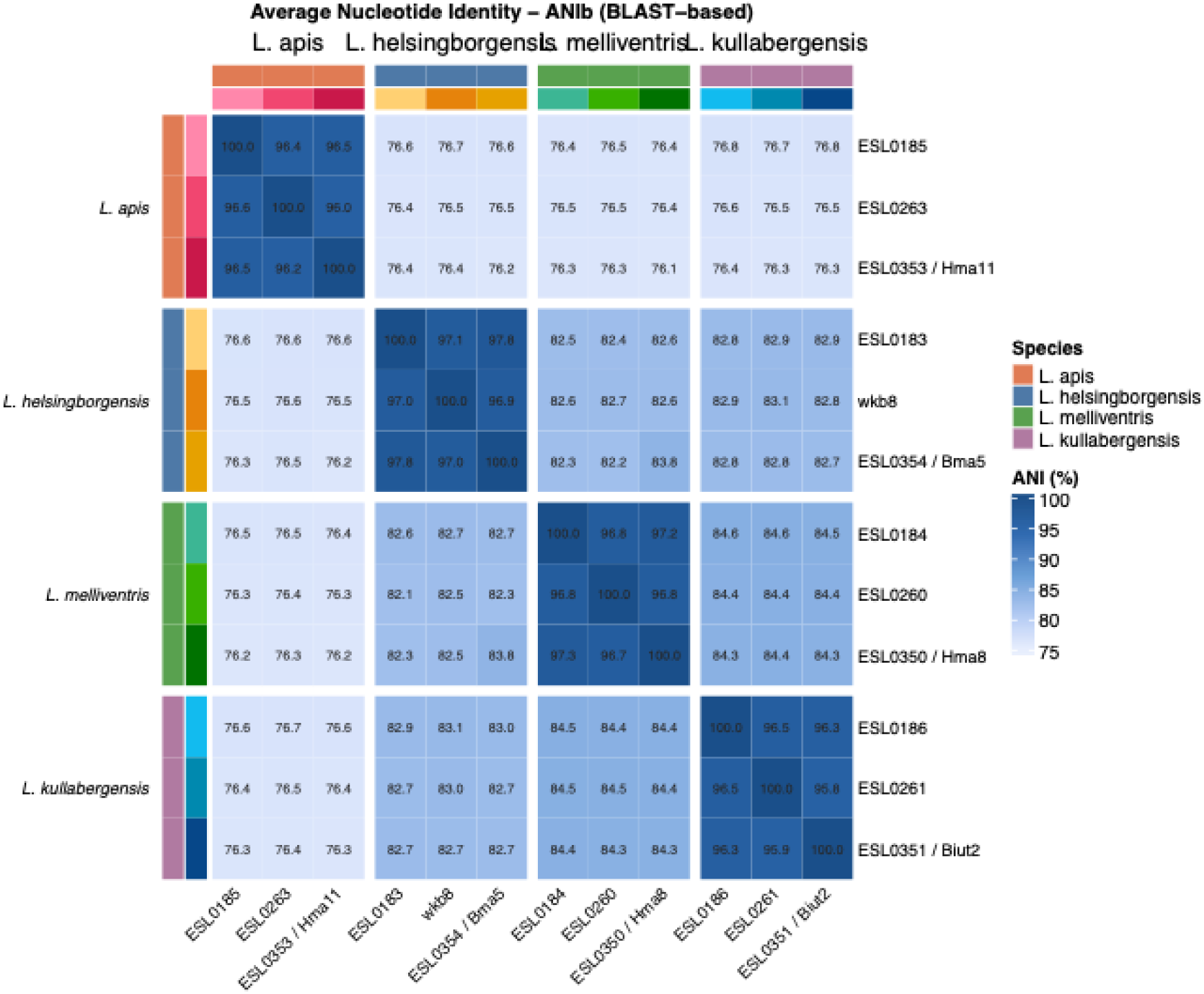
Pairwise average nucleotide identity (ANI) between the genomes of the 12 strains of this study. The strains of each of the four species are clustered together and indicated by strain name.

**Supplementary Figure 2.**
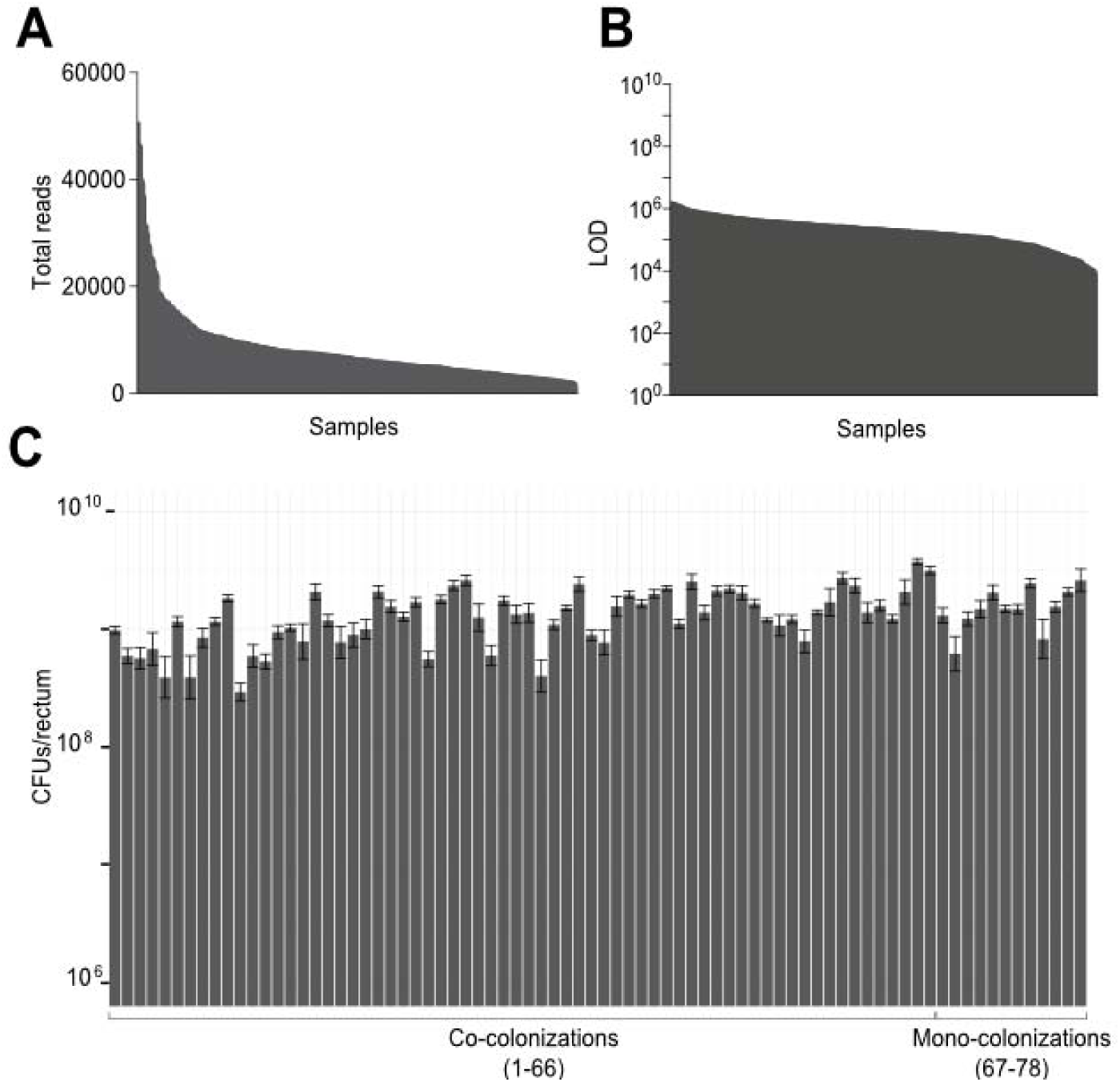
Total number of reads, limit of detection, and CFUs per rectum in each sample of the co-colonization experiment. **(A)** Total number of reads obtained for each of the 332 bee samples analyzed (12 mono-colonizations, 66 pairwise co-colonizations, all replicates included). **(B)** Limit of detection (LOD) calculated as the ratio of CFUs and total number of reads for each sample. **(C)** CFUs/rectum calculated for each sample (means of 3-5 bee samples, standard deviation from the mean is shown).

**Supplementary Figure 3.**
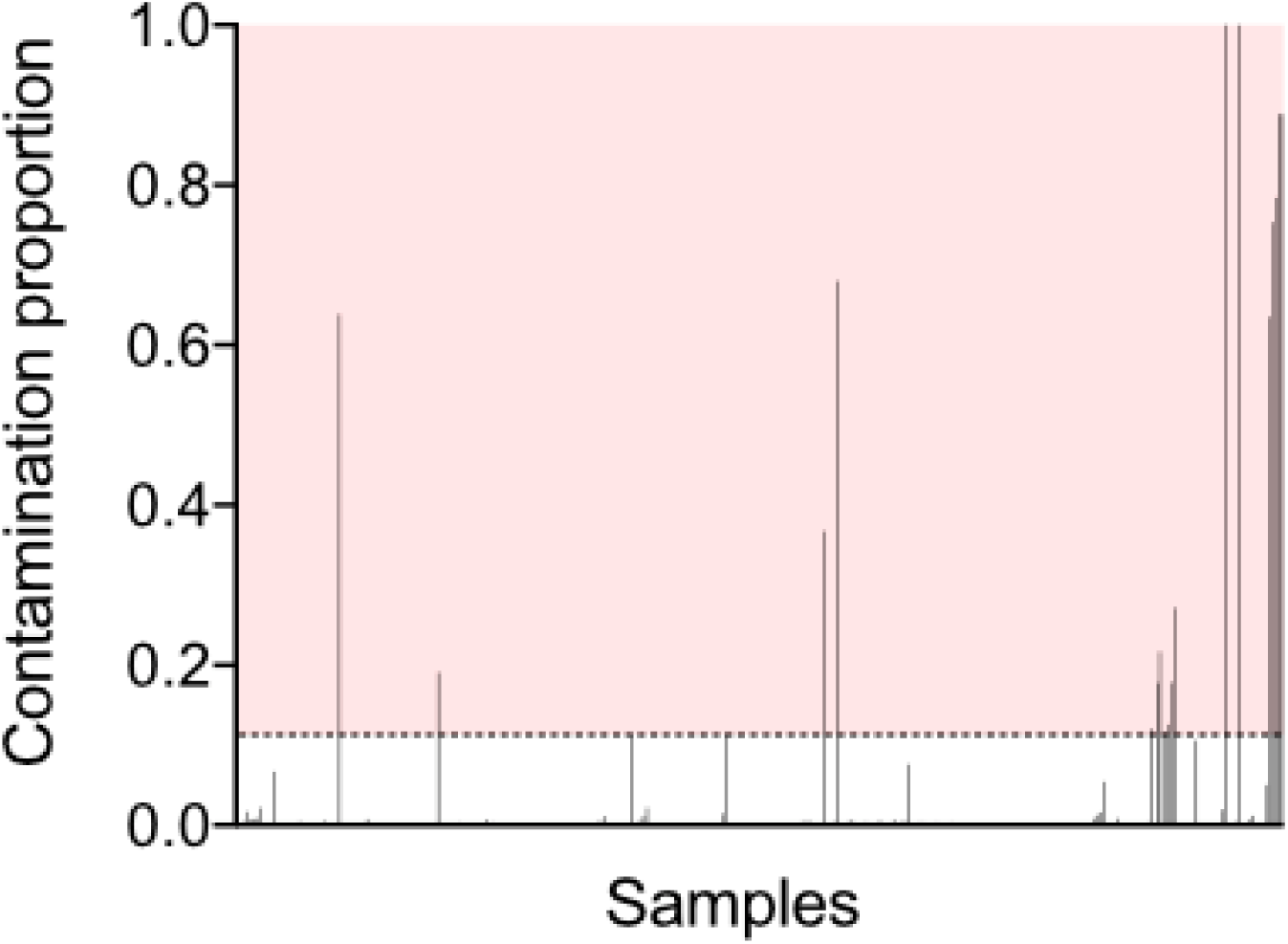
Identification of contaminations in each sample of the co-colonization experiment. The plot displays the proportion of reads assigned to a *mutT* gene variant that does not match the variant present in one of the strains in the inoculum in each sample. The light red area (contamination proportion > 10%) includes all the samples that were excluded for the downstream analysis.

**Supplementary Figure 4.**
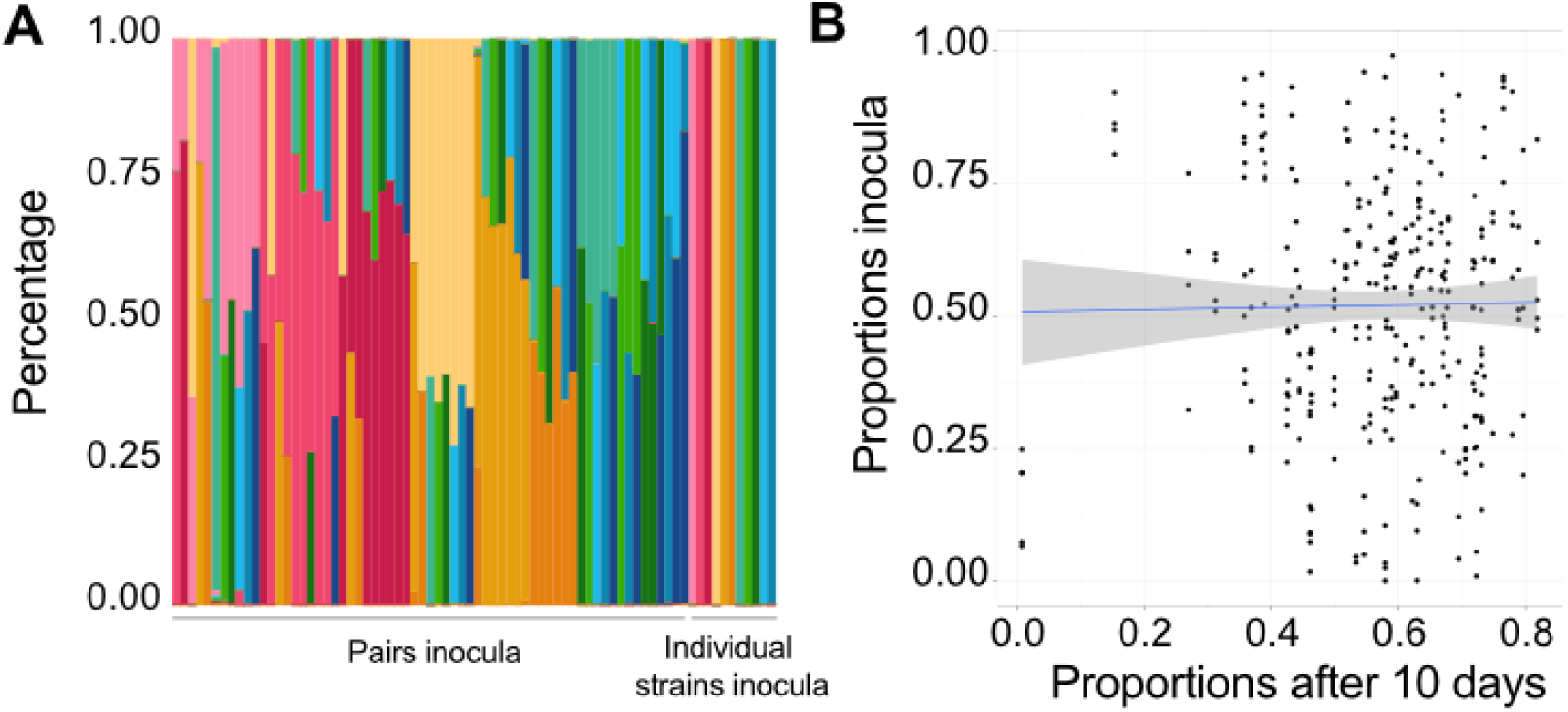
Strain proportions in the inocula and in gnotobiotic bees ten days after colonization across all samples of the co-colonization experiment. **(A)** Stacked bar plot of relative abundances in experimental inocula under paired- and single-strain conditions; colors indicate strains. (**B)** Correlation between strain proportions in the inoculum and at the end of the experiment (R² = 0.00022).

**Supplementary Figure 5.**
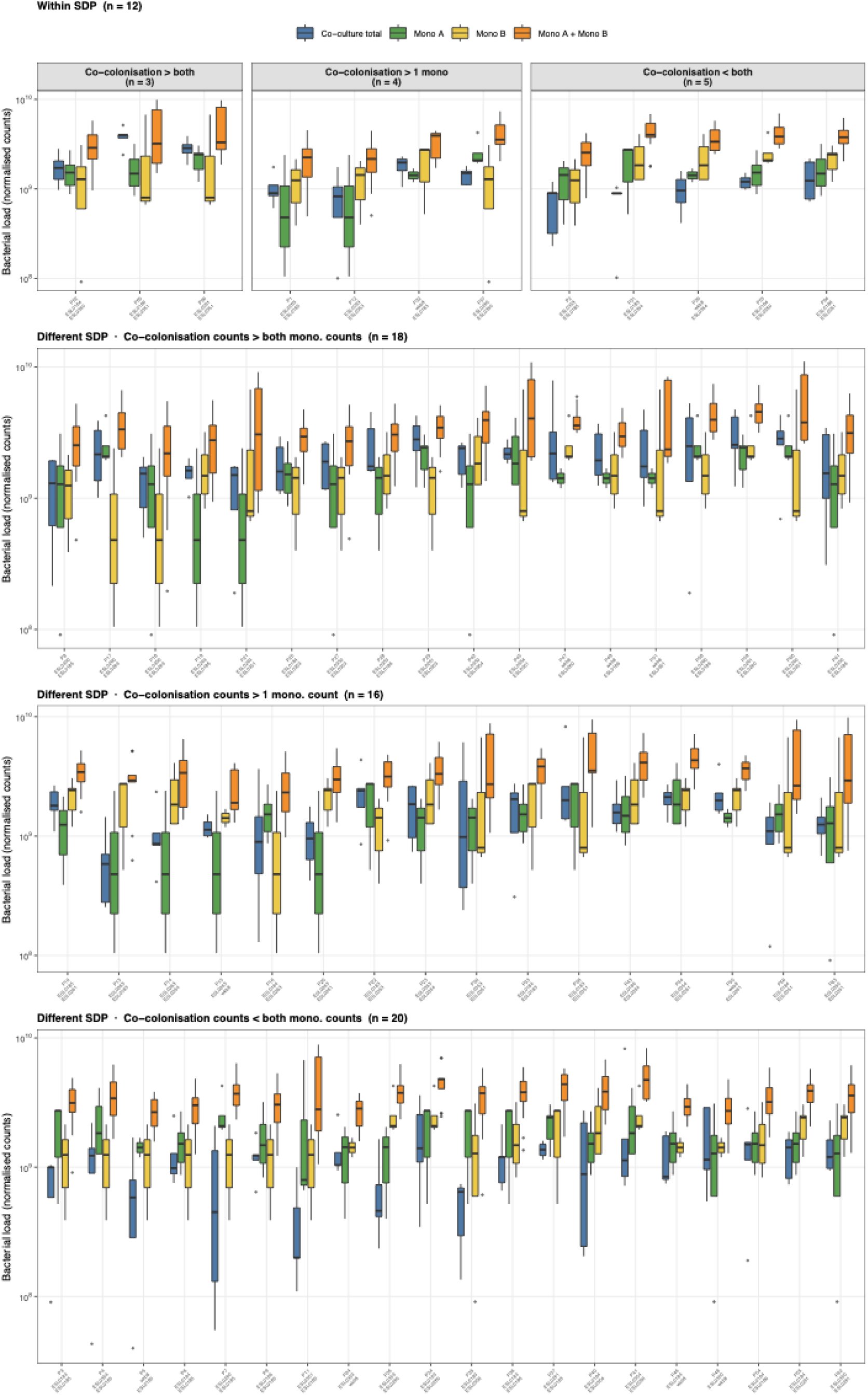
Total bacterial load in co-culture versus monoculture. For each strain pair (n = 66), boxplots show the distribution of total bacterial load (normalised CFU counts) across three conditions: co-culture (blue; sum of both strains), and each strain grown alone (Mono A, green; Mono B, yellow). The orange boxes show all pairwise sums of monoculture replicates (Mono A + Mono B), providing a null expectation under additive colonisation with no inter-strain interaction. Pairs are grouped by species identity (Within SDP vs. Different SDP) and further stratified by whether the median co-culture load exceeds both monoculture loads (“co > both”), only one (“co > 1 mono only”), or neither (“co < both”). The y-axis is log□□-scaled.

**Supplementary Figure 6.**
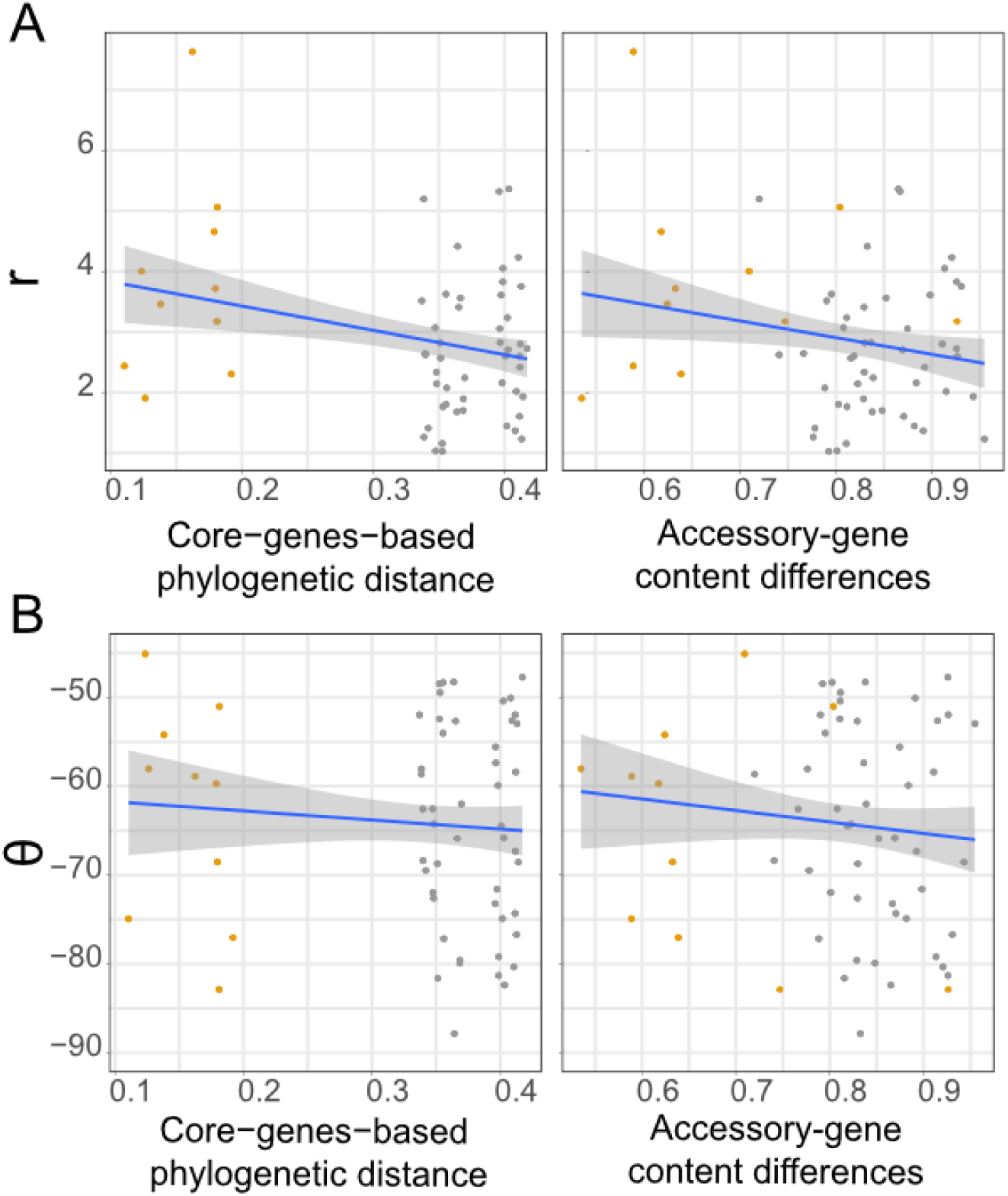
Correlation between interaction type and strength and phylogenetic distance and gene content differences across all tested strain pairs. **(A)** Correlation between each vector’s r (interaction-strength) and the 12 strains core-genes-based phylogenetic distance (R^2^ = 0.076) or accessory gene content differences (R^2^ = 0.043). Means of replicates (n=3-5) are displayed. **(B)** Correlation between each vector’s θ (interaction-type) and the 12 strains core-genes-based phylogenetic distance (R^2^ = 0.0062) or accessory gene content differences (R^2^ = 0.012). Means of replicates (n=3-5) are displayed. Dark yellow = conspecific strain-pairs; grey = allospecific strain-pairs.

**Supplementary Figure 7.**
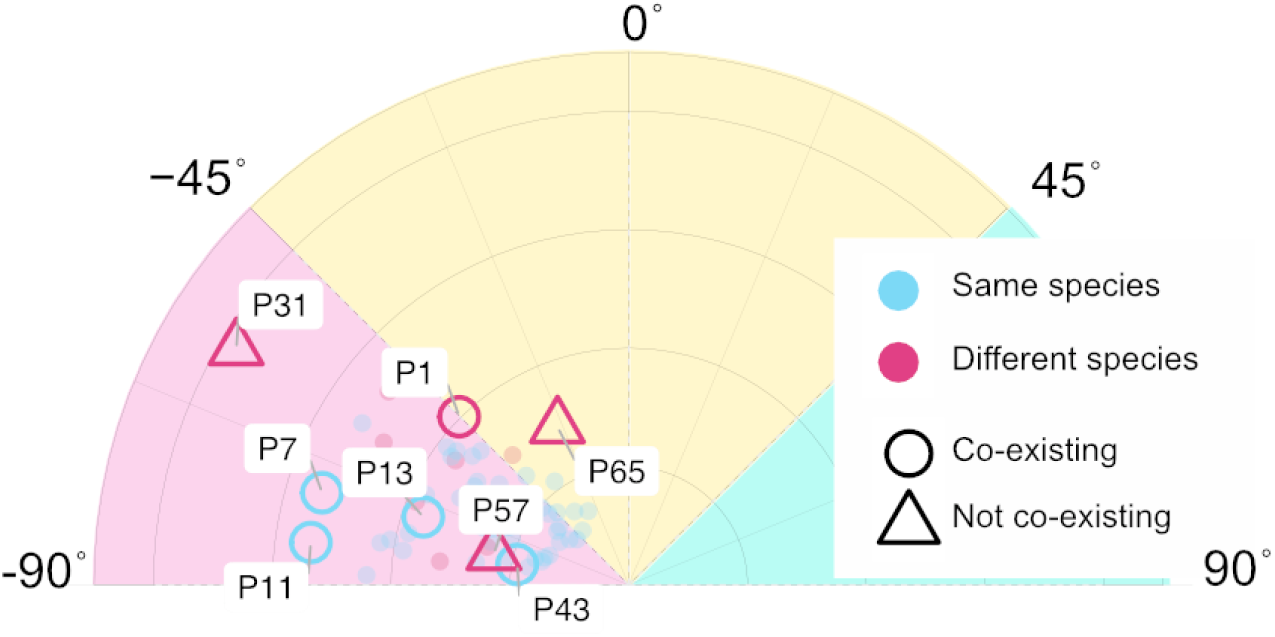
Polar plot highlighting the pairs randomly selected for the passaging experiment. Plot is the same as in Figure 2A. Colors and shape indicate within-vs between-species pairs and whether the two strains of these pairs coexisted across the three passages.

**Supplementary Figure 8.**
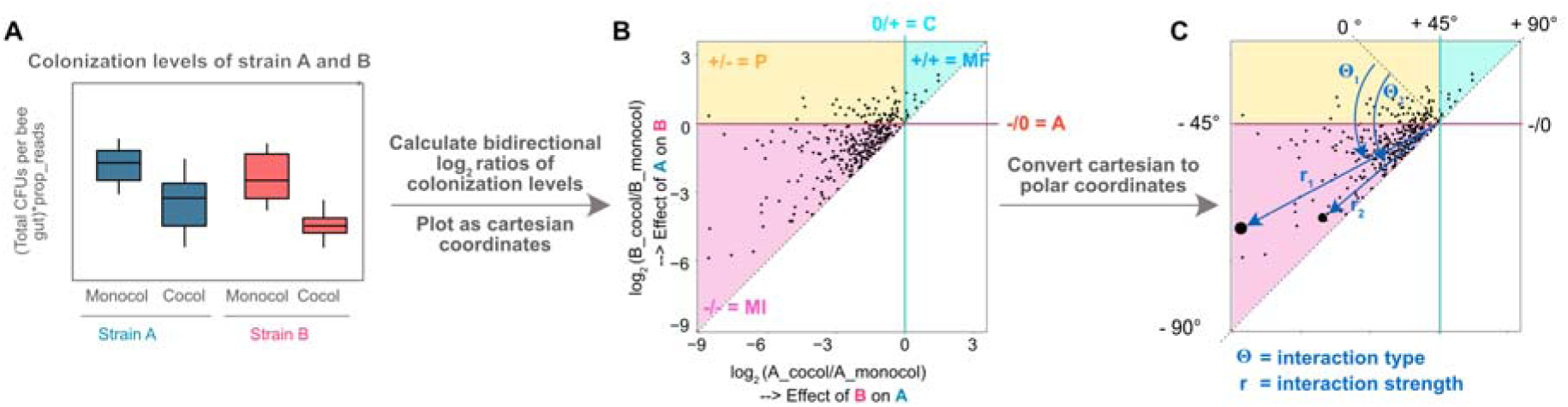
Framework to analyze the type and strength of pairwise interactions of *Lactobacillus* strains in the bee gut. **(A)** Schematic of colonization levels in co-colonization and mono-colonization for strains (A and B) in a given pair. The colonization level of each strain was calculated by multiplying the total number of CFUs by the proportion of sequencing reads assigned to that strain based on amplicon sequencing for each replicate. To quantify the effect of strain B on strain A, and vice versa, in each co-colonization replicate, we determined the log2 ratio of the colonization level of strain A in co-colonization and the mean colonization level of strain A in mono-colonization. (B) Cartesian coordinate plot of the reciprocal log2 ratios of the two strains in each replicate. The lower value of the two log2 ratios is always plotted on the x-axis so that all values appear above the diagonal. Depending on where a given strain pair falls its interaction type can be determined as indicated by different colors. 1/-= MI, mutual inhibition; -/0=A, amensamlism; +/-= P, parasitism;0/+= C, commensalism; +/+= MF, mutual facilitation. (C) Converting Cartesian coordinates to polar coordinates allows the reciprocal effects between each strain pair to be represented by an angle, θ, and a radius, r. The angle θ describes the interaction type, while the radius r represents the cumulative interaction strength between the two strains. In the two highlighted examples, an angle θ < −45° indicates mutual inhibition. However, θ□ is closer to −90° than θ□, indicating that the strains in pair 2 inhibit each other more symmetrically than those in pair 1. In contrast, the interaction strength, represented by r, is smaller for pair 2 than for pair 1. This means that although the inhibition in pair 2 is more balanced, the overall inhibitory effect between the two strains is weaker than in pair 1.

## References

1. Niche conservatism as an emerging principle in ecology and conservation biology.

2. Phylogenetic niche conservatism and the evolutionary basis of ecological speciation - Pyron - 2015 - Biological Reviews - Wiley Online Library. https://onlinelibrary.wiley.com/doi/full/10.1111/brv.12154. .

3. Malfertheiner L et al. Community conservatism is widespread across microbial phyla and environments. Nat Ecol Evol 2026;10:232–245. 10.1038/s41559-025-02957-4

4. Van Rossum T et al. Diversity within species: interpreting strains in microbiomes. Nature Reviews Microbiology 2020;18:491–506. 10.1038/s41579-020-0368-1

5. Ellegaard KM, Engel P. Beyond 16S rRNA community profiling: Intra-species diversity in the gut microbiota. Frontiers in Microbiology 2016;7:1–16. 10.3389/fmicb.2016.01475

6. Baud GLC et al. Turnover of strain-level diversity modulates functional traits in the honeybee gut microbiome between nurses and foragers. Genome Biology 2023;24:283. 10.1186/s13059-023-03131-4

7. Zhao S et al. Adaptive Evolution within Gut Microbiomes of Healthy People. Cell Host & Microbe 2019;25:656–667.e8. 10.1016/j.chom.2019.03.007

8. Truong DT et al. Microbial strain-level population structure and genetic diversity from metagenomes. Genome Research 2017;27:626–638. 10.1101/gr.216242.116

9. Garud NR et al. Evolutionary dynamics of bacteria in the gut microbiome within and across hosts. PLOS Biology 2019;17:e3000102. 10.1371/journal.pbio.3000102

10. Goyal A et al. Interactions between strains govern the eco-evolutionary dynamics of microbial communities. eLife 2022;11:e74987. 10.7554/eLife.74987

11. Cohan FM. Bacterial Species and Speciation. Syst Biol 2001;50:513–524. 10.1080/10635150118398

12. Van Rossum T et al. Diversity within species: interpreting strains in microbiomes. Nature Reviews Microbiology 2020;18:491–506. 10.1038/s41579-020-0368-1

13. HilleRisLambers J et al. Rethinking Community Assembly through the Lens of Coexistence Theory. *Annual Review of Ecology*, Evolution, and Systematics 2012;43:227–248. 10.1146/annurev-ecolsys-110411-160411

14. Orr JA, Armitage DW, Letten AD. Coexistence Theory for Microbial Ecology, and Vice Versa. Environmental Microbiology 2025;27:e70072. 10.1111/1462-2920.70072

15. Venkatanarayanan NN, Goyal A. Diverse communities promote the coexistence of closely-related strains through emergent equalization and stabilization. 2026;2026.03.16.711999. 10.64898/2026.03.16.711999

16. Biller SJ et al. Prochlorococcus: the structure and function of collective diversity. Nat Rev Microbiol 2015;13:13–27. 10.1038/nrmicro3378

17. Tettelin H et al. Genome analysis of multiple pathogenic isolates of Streptococcus agalactiae: Implications for the microbial “pan-genome”. Proceedings of the National Academy of Sciences 2005;102:13950–13955. 10.1073/pnas.0506758102

18. Welch RA et al. Extensive mosaic structure revealed by the complete genome sequence of uropathogenic Escherichia coli. Proceedings of the National Academy of Sciences 2002;99:17020–17024. 10.1073/pnas.252529799

19. Ellegaard KM, Engel P. Genomic diversity landscape of the honey bee gut microbiota. Nature Communications 2019;10.1:1–13.

20. Shoer S et al. Pangenomes of human gut microbiota uncover links between genetic diversity and stress response. Cell Host & Microbe 2024;32:1744–1757.e2. 10.1016/j.chom.2024.08.017

21. Hunt DE et al. Resource partitioning and sympatric differentiation among closely related bacterioplankton. Science 2008;320:1081–1085. 10.1126/science.1157890

22. Shapiro BJ et al. Population Genomics of Early Events in the Ecological Differentiation of Bacteria. Science 2012;336:48–51. 10.1126/science.1218198

23. Wolff R, Shoemaker W, Garud N. Ecological Stability Emerges at the Level of Strains in the Human Gut Microbiome. mBio 2023;14:e0250222. 10.1128/mbio.02502-22

24. Kramer J et al. Strain identity effects contribute more to Pseudomonas community functioning than strain interactions. ISME J 2025;19:wraf025. 10.1093/ismejo/wraf025

25. Brochet S et al. Niche partitioning facilitates coexistence of closely related honey bee gut bacteria. eLife 2021;10:e68583. 10.7554/eLife.68583

26. Yang C et al. Life history strategies complement niche partitioning to support the coexistence of closely related Gilliamella species in the bee gut. ISME J 2025;19:wraf016. 10.1093/ismejo/wraf016

27. Rakoff-Nahoum S, Foster KR, Comstock LE. The evolution of cooperation within the gut microbiota. Nature 2016;533:255–259. 10.1038/nature17626

28. Rakoff-Nahoum S, Coyne MJ, Comstock LE. An ecological network of polysaccharide utilization among human intestinal symbionts. Curr Biol 2014;24:40–49. 10.1016/j.cub.2013.10.077

29. Ratzke C, Barrere J, Gore J. Strength of species interactions determines biodiversity and stability in microbial communities. Nat Ecol Evol 2020;4:376–383. 10.1038/s41559-020-1099-4

30. Friedman J, Higgins LM, Gore J. Community structure follows simple assembly rules in microbial microcosms. Nat Ecol Evol 2017;1:0109. 10.1038/s41559-017-0109

31. Prasad A et al. Evolution of gut microbiota across honeybee species revealed by comparative metagenomics. Nature Communications 2025;16:9069. 10.1038/s41467-025-64115-5

32. Motta EVS, Moran NA. The honeybee microbiota and its impact on health and disease. Nat Rev Microbiol 2024;22:122–137. 10.1038/s41579-023-00990-3

33. Moran NA et al. Distinctive Gut Microbiota of Honey Bees Assessed Using Deep Sampling from Individual Worker Bees. PLoS ONE 2012;7:e36393. 10.1371/journal.pone.0036393

34. Engel P, Martinson VG, Moran NA. Functional diversity within the simple gut microbiota of the honey bee. Proceedings of the National Academy of Sciences 2012;109:11002–11007. 10.1073/pnas.1202970109

35. Zheng H et al. Honey bees as models for gut microbiota research. Lab Animal 2018;47:317–325. 10.1038/s41684-018-0173-x

36. Kešnerová L et al. Disentangling metabolic functions of bacteria in the honey bee gut. PLoS Biology 2017;15:1–51. 10.1371/journal.pbio.2003467

37. Ellegaard KM et al. Vast Differences in Strain-Level Diversity in the Gut Microbiota of Two Closely Related Honey Bee Species. Current Biology 2020;30:2520–2531.e7. 10.1016/j.cub.2020.04.070

38. Ellegaard KM et al. Genomic changes underlying host specialization in the bee gut symbiont Lactobacillus Firm5. Molecular Ecology 2019;28:2224–2237.

39. Kehe J et al. Positive interactions are common among culturable bacteria. Sci Adv 2021;7159:1–11. 10.1101/2020.06.24.169474

40. Pycke JR. Some tests for uniformity of circular distributions powerful against multimodal alternatives. Canadian Journal of Statistics 2010;38:80–96. 10.1002/cjs.10048

41. Arevalo P et al. A Reverse Ecology Approach Based on a Biological Definition of Microbial Populations. Cell 2019;178:820–834.e14. 10.1016/j.cell.2019.06.033

42. Shapiro BJ, Polz MF. Microbial Speciation. Cold Spring Harbor Perspectives in Biology 2015;7:a018143. 10.1101/cshperspect.a018143

43. Conrad RE et al. Microbial species and intraspecies units exist and are maintained by ecological cohesiveness coupled to high homologous recombination. Nat Commun 2024;15:9906. 10.1038/s41467-024-53787-0

44. Martinson VG et al. A simple and distinctive microbiota associated with honey bees and bumble bees. Molecular Ecology 2011;20:619–628. 10.1111/j.1365-294X.2010.04959.x

45. Poyet M et al. A library of human gut bacterial isolates paired with longitudinal multiomics data enables mechanistic microbiome research. Nat Med 2019;25:1442–1452. 10.1038/s41591-019-0559-3

46. Lloyd-Price J et al. Strains, functions and dynamics in the expanded Human Microbiome Project. Nature 2017;550:61–66. 10.1038/nature23889

47. Strachan CR et al. Differential carbon utilization enables co-existence of recently speciated Campylobacteraceae in the cow rumen epithelial microbiome. Nature Microbiology 2023;8:309–320. 10.1038/s41564-022-01300-y

48. Bobay L-M, Wissel EF, Raymann K. Strain Structure and Dynamics Revealed by Targeted Deep Sequencing of the Honey Bee Gut Microbiome. mSphere 2020;5:10.1128/msphere.00694-20. 10.1128/msphere.00694-20

49. Powell E, Ratnayeke N, Moran NA. Strain diversity and host specificity in a specialized gut symbiont of honeybees and bumblebees. Molecular Ecology 2016;25:4461–4471. 10.1111/mec.13787

50. Leventhal GE et al. Strain-level diversity drives alternative community types in millimetre-scale granular biofilms. Nat Microbiol 2018;3:1295–1303. 10.1038/s41564-018-0242-3

51. Zhang C, Zhao L. Strain-level dissection of the contribution of the gut microbiome to human metabolic disease. Genome Med 2016;8:41. 10.1186/s13073-016-0304-1

52. Yan Y et al. Strain-level epidemiology of microbial communities and the human microbiome. Genome Medicine 2020;12:71. 10.1186/s13073-020-00765-y

53. Prasad A et al. Priority effects drive strain-level community composition of honeybee gut microbiota. ISME J 2026;20:wrag056. 10.1093/ismejo/wrag056

54. Jones KR et al. Effects of priority on strain-level composition of the honey bee gut community. Appl Environ Microbiol 2025;91:e00828–25. 10.1128/aem.00828-25

55. Loreau M, Hector A. Partitioning selection and complementarity in biodiversity experiments. Nature 2001;412:72–76. 10.1038/35083573

56. Kešnerová L et al. Gut microbiota structure differs between honeybees in winter and summer. The ISME Journal 2020;14:801–814. 10.1038/s41396-019-0568-8

57. Kešnerová L et al. Gut microbiota structure differs between honeybees in winter and summer. ISME J 2020;14:801–814. 10.1038/s41396-019-0568-8

58. Russel J et al. Antagonism correlates with metabolic similarity in diverse bacteria. Proceedings of the National Academy of Sciences 2017;114:10684–10688. 10.1073/pnas.1706016114

59. Ellegaard KM et al. Genomic changes underlying host specialization in the bee gut symbiont Lactobacillus Firm5. Molecular Ecology 2019;28:2224–2237. 10.1111/mec.15075

60. Picot A et al. Microbial interactions in theory and practice: when are measurements compatible with models? Current Opinion in Microbiology 2023;75:102354. 10.1016/j.mib.2023.102354

61. Adler PB et al. What Have We Learned From Empirical Applications of Modern Coexistence Theory? Ecology Letters 2026;29:e70404. 10.1111/ele.70404

62. Chesson P. Mechanisms of Maintenance of Species Diversity. Annual Review of Ecology, Evolution, and Systematics 2000;31:343–366. 10.1146/annurev.ecolsys.31.1.343

63. Barabás G, D’Andrea R, Stump SM. Chesson’s coexistence theory. Ecological Monographs 2018;88:277–303. 10.1002/ecm.1302

64. McAvoy T et al. Sign and strength of pairwise interactions in natural isolates depend on environment type. 2026;2026.03.31.715556. 10.64898/2026.03.31.715556

65. Sundarraman D et al. Higher-Order Interactions Dampen Pairwise Competition in the Zebrafish Gut Microbiome. mBio 2020;11:10.1128/mbio.01667-20. 10.1128/mbio.01667-20

66. Ishizawa H et al. Learning beyond-pairwise interactions enables the bottom–up prediction of microbial community structure. Proceedings of the National Academy of Sciences 2024;121:e2312396121. 10.1073/pnas.2312396121

67. Baichman-Kass A, Song T, Friedman J. Competitive interactions between culturable bacteria are highly non-additive. eLife 2023;12:e83398. 10.7554/eLife.83398

68. O’ Donnell MM et al. Carbohydrate catabolic flexibility in the mammalian intestinal commensal Lactobacillus ruminis revealed by fermentation studies aligned to genome annotations. Microb Cell Fact 2011;10:S12. 10.1186/1475-2859-10-S1-S12

69. Bolger AM, Lohse M, Usadel B. Trimmomatic: a flexible trimmer for Illumina sequence data. Bioinformatics 2014;30:2114–2120. 10.1093/bioinformatics/btu170

70. Zhang J et al. PEAR: a fast and accurate Illumina Paired-End reAd mergeR. Bioinformatics 2014;30:614–620. 10.1093/bioinformatics/btt593

